# Integrated multi-omics reveals adaptive anti-oxidant remodeling in early alcohol-associated liver disease

**DOI:** 10.64898/2026.05.28.728507

**Authors:** Tsung-Heng Tsai, Mirjavid Aghayev, Serguei Ilchenko, Usman Sabir, Megan R. McMullen, Salan Ghaju, Xiaoxin Luke Chen, Guo-Fang Zhang, Wilson C.J. Chung, Laura E. Nagy, Takhar Kasumov

**Affiliations:** Department of Mathematical Sciences, Kent State University, Kent, OH, 44242; Department of Pharmaceutical Sciences, College of Pharmacy, Northeast Ohio Medical University, Rootstown, OH, 44272; Departments of Inflammation and Immunity and Gastroenterology/Hepatology, Northern Ohio Alcohol Center, Cleveland Clinic, Cleveland, OH, 44195; School of Biomedical Sciences, Kent State University, Kent, OH, 44242; Coriell Institute for Medical Research, 403 Haddon Avenue, Camden, NJ, 08103; Department of Surgery, Cooper University Health Care, 401 Haddon Avenue, Camden, NJ, 08103; Division of Division of Endocrinology, Metabolism and Nutrition, Duke Molecular Physiology Institute, and Department of Medicine, Duke University, Durham, NC, 27701; Department of Biological Sciences, Kent State University, Kent, OH, 44242

## Abstract

Alcohol-associated liver disease (ALD) is a leading cause of liver-related morbidity and mortality. Although various omics approaches have revealed early metabolic alterations, individual datasets provide limited mechanistic insight. Here, we integrated RNA sequencing with mass spectrometry–based analyses to quantify gene expression, protein abundance, proteome and acetylome dynamics, and metabolic fluxes in livers of alcohol-fed mice. This multi-layered approach revealed extensive metabolic rewiring characterized by suppressed mitochondrial energy metabolism and compensatory upregulation of glutathione (GSH) production, utilization, and recycling, establishing a high-flux antioxidant network. These changes were coupled to epigenetic histone H3 remodeling, marked by increased permissive acetylation and decreased suppressive methylation, linking alcohol-induced metabolic and redox alterations to chromatin reprogramming. ChEA-based *in silico* upstream transcription factor analysis, identified hepatocyte nuclear factor 4α (HNF4α) and nuclear factor erythroid 2–related factor 2 (NRF2) as key regulatory nodes. Alcohol exposure was associated with a modest HNF4α suppression alongside increased expression of NRF2, indicating a shift from HNF4α-driven metabolic programs toward NRF2-mediated antioxidant responses. Despite acetylation-associated impairment of mitochondrial proteins, GSH-related enzymes were preserved, supporting a protective, high-turnover antioxidant response that limits early oxidative stress and defines an adaptive state maintaining redox homeostasis while potentially predisposing to ALD progression.

## INTRODUCTION

The liver is a central hub of metabolic homeostasis, coordinating diverse physiological processes essential for systemic health, including regulation of glucose and lipid metabolism, ammonia detoxification, bile synthesis, and xenobiotic clearance. As the primary site of ethanol metabolism, the liver is vulnerable to alcohol-induced injury, a key driver in the pathogenesis of alcohol-associated liver disease (ALD)^1^. Chronic alcohol consumption promotes hepatic injury through the generation of toxic metabolites, including acetaldehyde and reactive oxygen species (ROS), which impair mitochondrial function, disrupt redox balance, and alter intermediary metabolism.

In the early stages of ALD, hepatic steatosis develops in the presence of relatively modest inflammation and oxidative damage, suggesting engagement of adaptive or compensatory mechanisms that transiently preserve cellular homeostasis^2^. Nonetheless, alcohol-induced perturbations in lipid metabolism promote hepatic lipid accumulation and lipotoxicity, accompanied by increased lipid peroxidation and oxidative stress^3^. Mitochondria are central to this process, acting as both a source and target of ROS^1^. Chronic alcohol exposure impairs electron transport chain (ETC) function, particularly at Complexes I and III, increasing electron leakage and superoxide generation^4^. Resulting oxidative damage to mitochondrial DNA, proteins, and membrane lipids compromises mitochondrial function^5^. These processes place sustained demand on hepatic antioxidant defenses^1^, among which glutathione (GSH) serves as the principal intracellular redox buffer and a critical regulator of mitochondrial integrity and cell survival^6^. While depletion of hepatic GSH has been linked to advanced stages of ALD, accumulating evidence indicates that early alcohol exposure may instead activate compensatory antioxidant responses aimed at maintaining redox balance^7^.

Beyond transcriptional regulation, alcohol exposure induces extensive remodeling of the hepatic proteome, altering protein abundance, stability, and post-translational modification (PTM) landscapes in ways that are not fully captured by mRNA-based analyses alone^8^. Proteomic studies have shown that chronic alcohol consumption reshapes proteins involved in mitochondrial metabolism, lipid synthesis, oxidation and storage, antioxidant defense, and proteostasis, thereby altering metabolic capacity at the post-transcriptional level^9,10^. Among these regulatory layers, lysine acetylation has emerged as a prominent alcohol-responsive modification, reflecting changes in acetyl-CoA availability, mitochondrial–cytosolic metabolic coupling, and cellular redox state^11^. Alcohol metabolism not only induces oxidative stress but also elevates acetyl-CoA levels while reducing NAD⁺ availability through an increased NADH/NAD⁺ ratio^12^. Together, these shifts enhance acetyltransferase activity and constrain NAD⁺-dependent deacetylases (e.g., sirtuins), thereby promoting global protein hyperacetylation.

Protein acetylation has been recognized as a dynamic regulator of protein function and stability, influencing enzymatic activity, subcellular localization, and protein degradation pathways^11^. Alcohol-induced hyperacetylation of hepatic proteins has been associated with impaired mitochondrial metabolism, altered fatty acid oxidation, and dysregulated stress responses. Importantly, a growing body of evidence suggests that lysine acetylation directly modulate protein function and turnover, providing a mechanistic link through which metabolic state impacts enzyme activity and proteome dynamics^10,13,14^. Recent proximity proteomics studies further demonstrated that alcohol-induced lysine acetylation preferentially occurs within cysteine–lysine motifs, where cysteine-proximal acetylation is associated with redox remodeling of the hepatic proteome, particularly affecting pathways involved in mitochondrial bioenergetics and GSH metabolism that are disrupted in ALD^15,16^. Together, these findings underscore the importance of integrated multi-omics approaches for defining the complex interplay among lysine acetylation, redox signaling, and metabolic dysfunction in ALD.

Studies integrating multi-omics datasets have enabled a more comprehensive characterization of ALD pathogenesis^17^. For example, an integrative analysis combining genome-wide association studies (GWAS), DNA methylation, and RNA sequencing (RNA-seq) demonstrated that ALD progression is associated with abnormal transcriptomic reprogramming, including defective activity of liver-enriched transcription factors such as hepatocyte nuclear factor 4α (HNF4α) caused by altered DNA methylation and chromatin remodeling^18^. Complementary approaches integrating metabolomic and transcriptomic data have identified key metabolic bottlenecks in alcohol-exposed liver, including the accumulation of glucose-6-phosphate and induction of hexokinase domain containing 1, the latter emerging as a strong predictor of patient survival^19^. Studies combining transcriptomic and proteomic profiling have further facilitated the identification of candidate plasma biomarkers in ALD^9,20^, highlighting the translational potential of such multi-layered approaches.

While these traditional static multi-omics approaches have significantly advanced our understanding of ALD pathogenesis and enabled noninvasive characterization of distinct disease stages, they offer only temporal snapshots of the transcriptome, proteome, and metabolome. As such, steady-state measurements often fail to capture dynamic regulatory processes, including protein turnover and PTMs, that underlie cellular adaptation. To address this limitation, we integrated static multi-omics with quantitative analyses of proteome and acetylome dynamics to define early adaptive responses to chronic ethanol exposure in mice. By coupling these datasets with *in silico* upstream regulator analysis, this approach revealed perturbations in mitochondrial bioenergetics. These alterations appear to trigger a compensatory activation of antioxidant pathways, particularly GSH homeostasis. To directly quantify metabolic activity beyond inferred pathways, we applied stable isotope–resolved flux analysis using ²H₂O metabolic labeling to quantify hepatic reduced (GSH) and oxidized glutathione (GSSG) turnover, including synthesis, utilization, and recycling rates. This kinetic approach provides a high-resolution view of how the liver dynamically maintains redox balance under sustained ethanol-induced oxidative stress. Together with widespread changes in protein turnover, these findings reveal a coordinated adaptive response linking redox regulation, metabolic remodeling, and proteome stability during early ALD pathogenesis.

## RESULTS

We generated RNA-seq (n=12), label-free quantification (LFQ) proteomics (n=18), and kinetic proteomics data from a collection of 18 female mouse liver tissues: 9 pair-fed (PF) and 9 ethanol-fed (EF). The quantitative and kinetic proteomics, together with relevant targeted metabolomics data have been published^10^. In that study, the analysis focused on proteins and acetylated sites with differential abundance or turnover rate, and their associated metabolic consequences in substrate metabolism under PF and EF conditions.

Here, we integrate newly generated RNA-seq data with previously published proteomics, kinetics of native and acetylated proteins, and metabolomics datasets to obtain a more comprehensive, system-level understanding of metabolic remodeling in ALD. We further interrogated a central finding of these analyses, compensatory activation of antioxidant pathways in response to alcohol-induced mitochondrial perturbations, particularly glutathione homeostasis, using stable isotope–resolved flux analysis. A schematic summary of the study design and analysis is provided in **Figure 1**.

**Figure 1.**
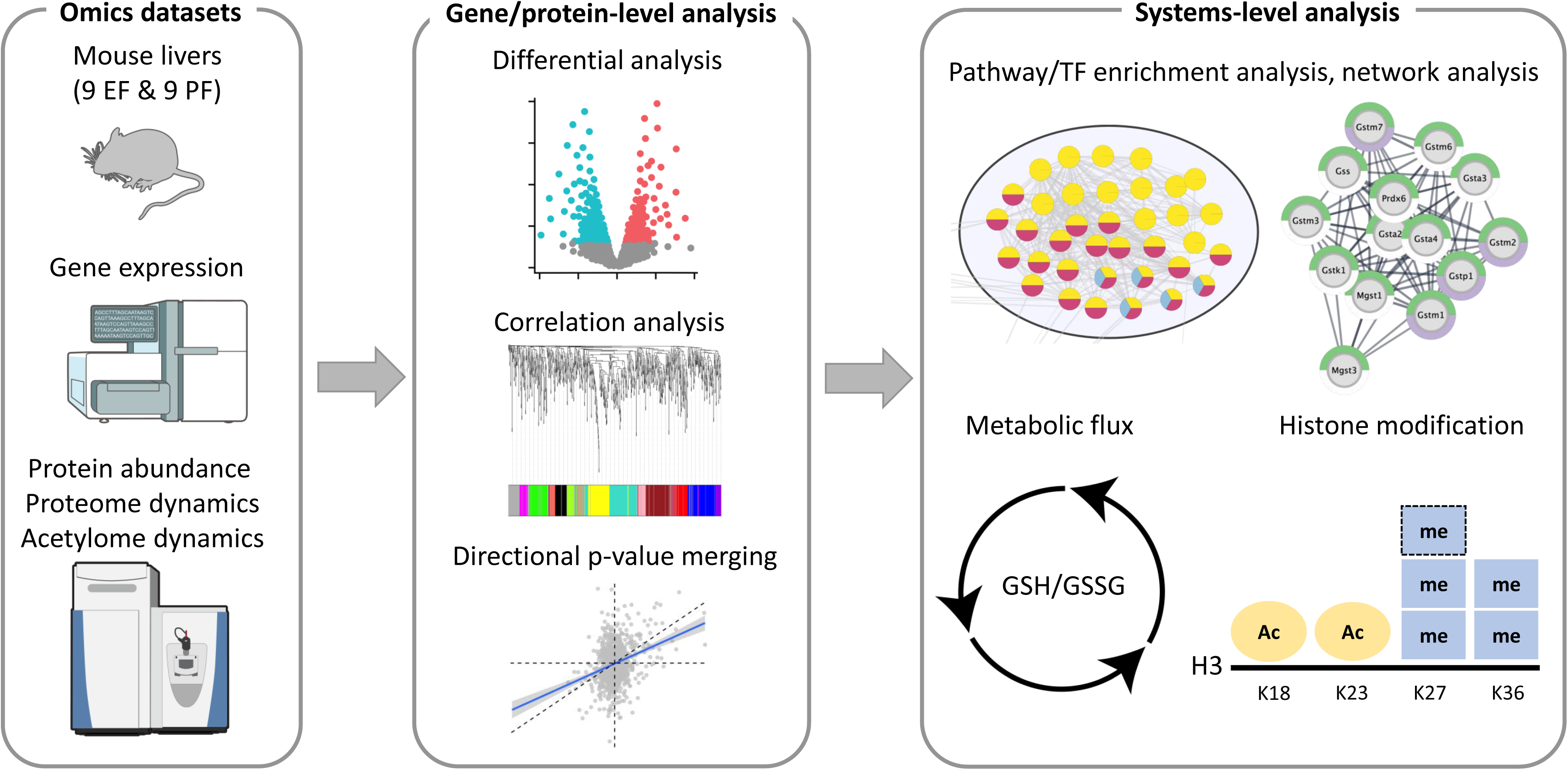
Study design and analytical workflow for integrated omics analysis of alcoholic liver disease in mice.

### Individual analyses of RNA-seq and LFQ proteomics datasets

The RNA-seq analysis quantified 17,003 mRNAs in 12 mouse livers (6 EF vs 6 PF), which identified a total of 535 differentially expressed genes (DEGs): 288 up-regulated and 247 down-regulated mRNAs in EF mice compared to PF, with an adjusted p<0.05 and at least two-fold change (i.e., |log_2_FC|>1) (**Figure 2A**, **Supplementary Table S1A**). The two groups of mice were clearly separated based on these differentially expressed mRNAs (**Figure 2B**). Using Gene Ontology Biological Process (GO:BP) terms as reference, we performed enrichment analysis with the DEGs. The analysis identified 212 and 25 enriched biological processes, respectively, associated with the up-regulated and down-regulated mRNAs (**Supplementary Table S1B**). Overall, several metabolic processes, including xenobiotic, fatty acid, retinoic acid, olefinic compound and icosanoid, emerged to be significantly enriched in both the up-regulated and down-regulated genes (**Figure 2C**-**D**). Additionally, genes up-regulated in EF were uniquely enriched in several biological processes related to hepatic stress responses to toxic substances and oxidative stress, with antioxidant glutathione metabolism prominently represented among the top-ranked pathways, alongside sulfur compound and vitamin metabolism. In contrast, genes down-regulated in EF were primarily associated with bile acid and bile salt transport and steroid metabolic processes (**Figure 2C**). Disruption of bile acid transport may contribute to hepatocellular injury, as bile acids regulate hepatic sulfur amino acid metabolism and glutathione-dependent antioxidant defenses, thereby influencing the liver’s susceptibility to oxidative stress^21^.

**Figure 2.**
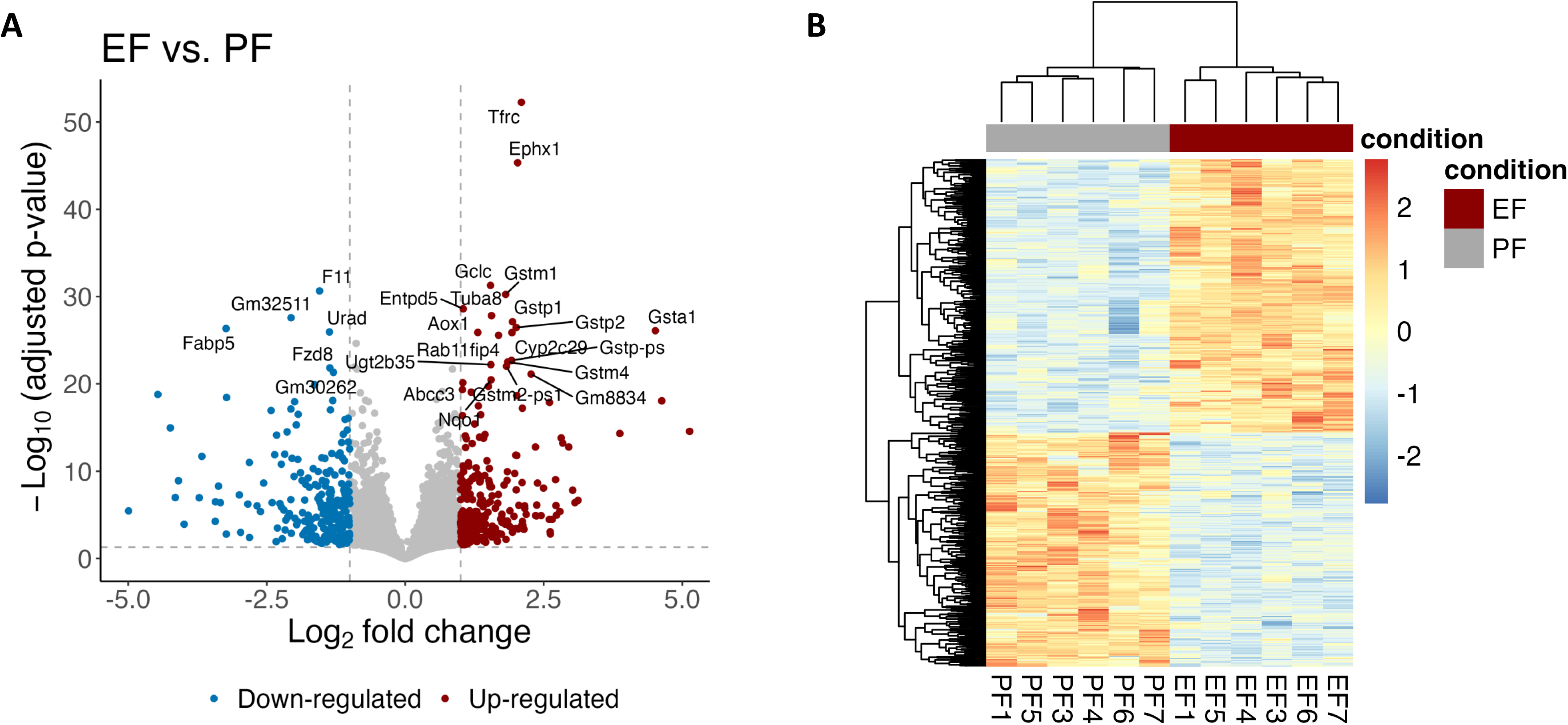
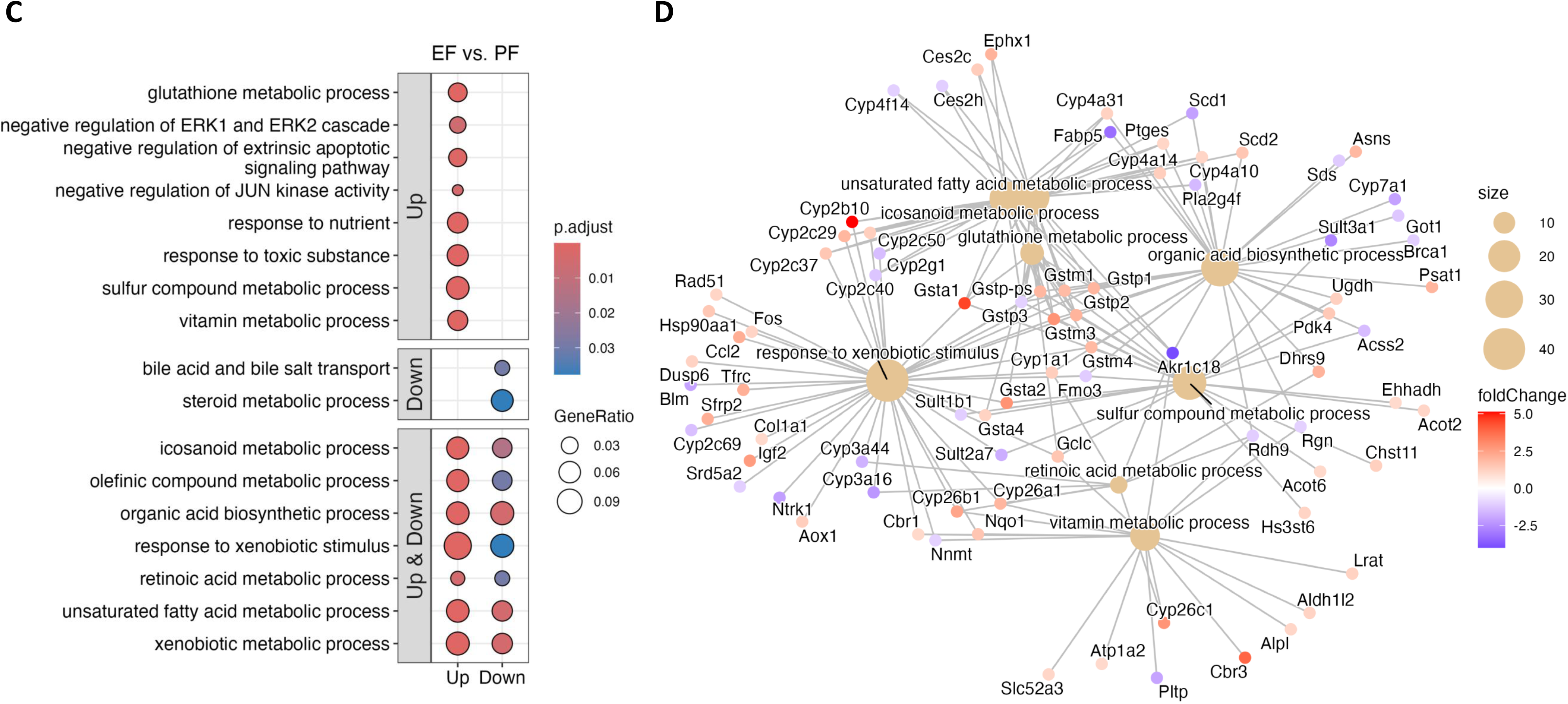
RNA-seq analysis of EF and PF mouse livers. (**A**) Differential gene expression analysis using DESeq2. Horizontal dashed line indicates the specified threshold for statistical significance (adjusted p-value of 0.05); vertical dashed lines represent two-fold increase or decrease of gene expression. (**B**) Heatmap of differentially expressed genes (adjusted p-value <0.05, |log_2_FC| >1) in the samples. (**C**) Representative enriched GO biological processes associated with up– and down-regulated mRNAs in EF vs. PF at the mRNA level. The enriched GO terms are organized in three panels to distinguish the terms identified with up-regulated mRNAs only, down-regulated mRNAs only, and both up– and down-regulated mRNAs. For example, glutathione process is associated with up-regulated mRNAs in EF. Full list of the enriched GO terms is provided in **Supplementary Table S1B**. (**D**) Gene network for the top significant GO terms (brown dots) and their associated genes with color-coded fold changes (red: up-regulated, blue: down-regulated).

In addition to the RNA-seq analysis, reanalysis of our previously published LFQ proteomics dataset^10^ identified alcohol-responsive proteins enriched in pathways related to fatty acid metabolism, xenobiotic responses, protein folding, and metabolic adaptation (**Supplementary Figure S1**, **Supplementary Table S2**). Co-expression network analysis of RNA-seq and proteomics datasets further identified alcohol-associated mRNA and protein modules linked to cellular catabolic, amino acid, and mitochondrial metabolic processes, supporting coordinated transcriptomic and proteomic remodeling in ALD (**Supplementary Figure S2**, **Supplementary Table S3**). However, the absence of significant cross-omics enrichment indicated that these changes were represented by distinct and weakly overlapping co-expression structures at the mRNA and protein levels. We therefore next sought to integrate mRNA and protein expression evidence in a more systematic and coherent manner.

### Integrating mRNA and protein data reveals early metabolic shifts in ALD that single-omics analyses overlook

Using the RNA-seq and LFQ proteomics data, we performed an integrated analysis combining the evidence at the mRNA and protein levels about the 1069 genes quantified by both omics analyses (**Figure 3A**, **Supplementary Table S4A**). Out of these genes, 551 and 167 had a p-value <0.05 in the RNA-seq and proteomics analyses, respectively, and 577 had the same direction of changes (EF vs. PF) in both datasets.

**Figure 3.**
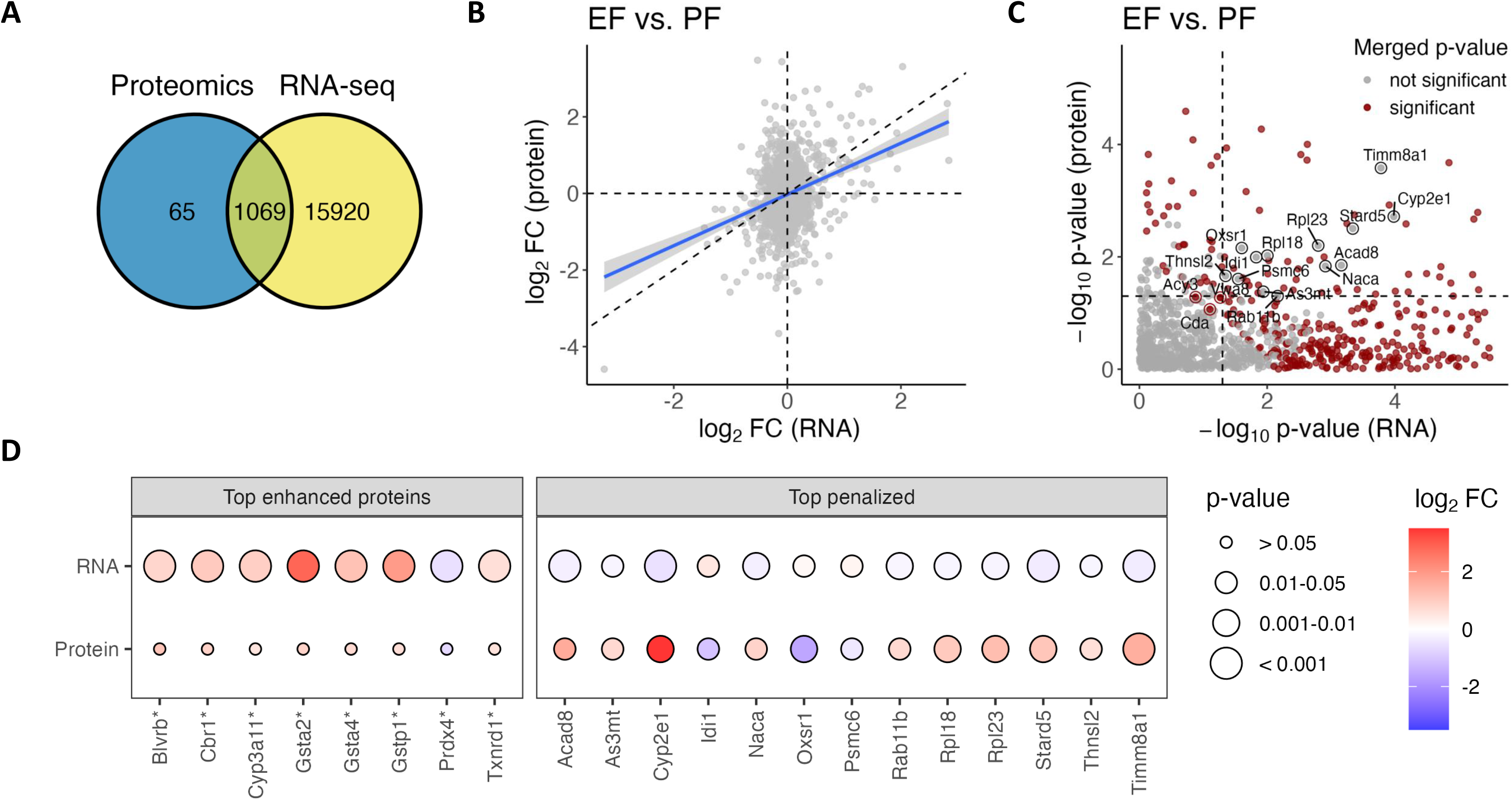
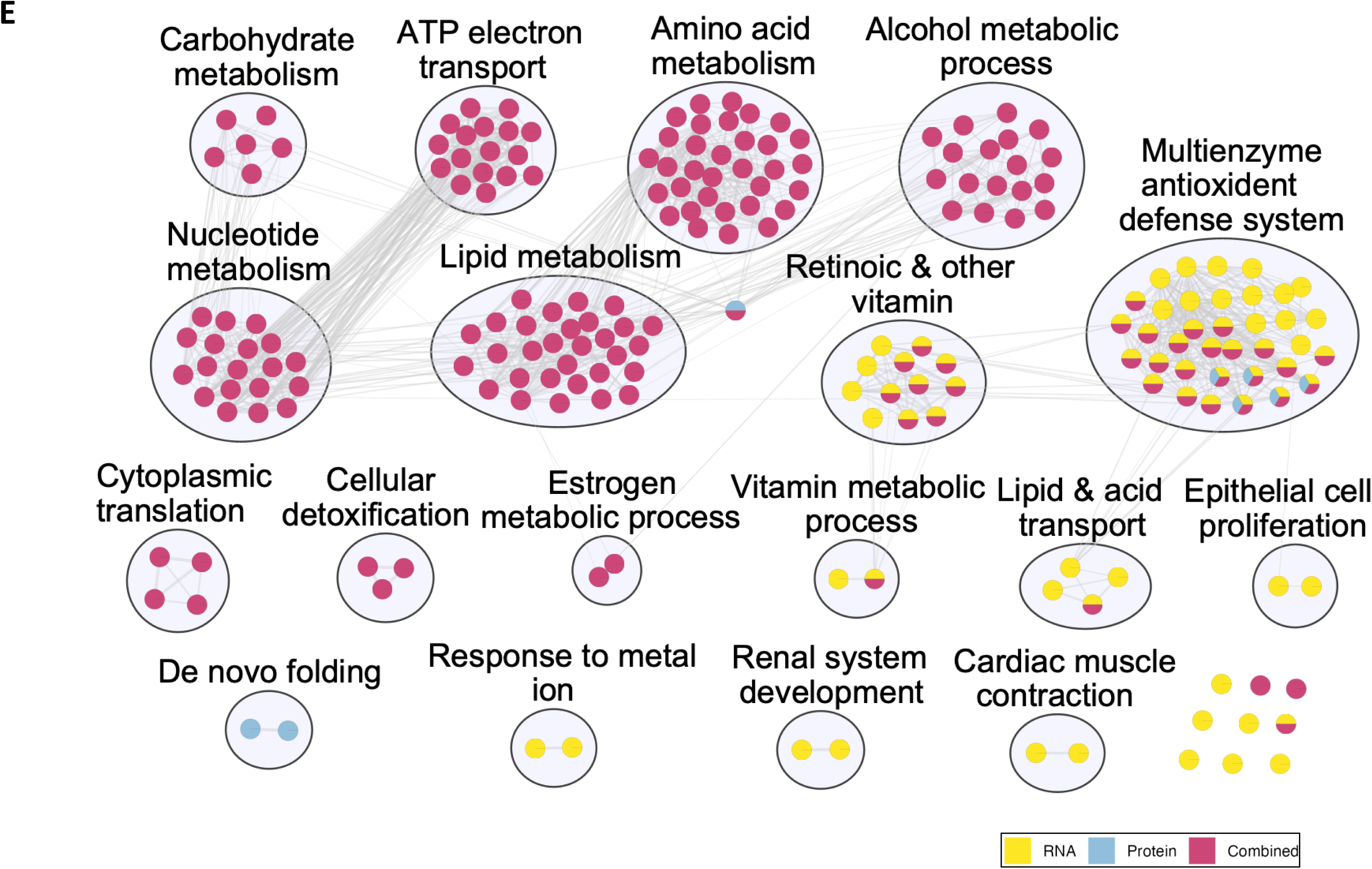
Directional p-value merging based on the fold changes and p-values at the mRNA and protein levels. (**A**) Number of quantified genes at the mRNA and protein levels. There are 1069 genes quantified in both RNA-seq and label-free proteomics analyses. (**B**) Correlation of changes at the mRNA and protein levels (on log_2_ scale) for the 1069 genes. (**C**) Statistical significance determined based on the mRNA (x-axis, more significant rightward) and protein (y-axis, more significant upward) levels, and their combined evidence (red: significant) using directional p-value merging implemented in ActivePathways^22^. Penalized (significant in both, non-significant after p-value merging, due to disagreement in fold change direction) and prioritized (not significant in both, but significant after combination) genes are highlighted. The plot is cropped, where 129 genes with an extremely small p-value at the mRNA level are not included. (**D**) Fold changes and p-values of example genes by the directional p-value merging. Top penalized genes are annotated as circled gray points in (**C**). Top enhanced proteins are genes/proteins that were not significant at the protein level but identified as significant after combining the mRNA information with consistent FC direction. These genes, including Gsta2, Gsta4 and Gstp1 involved in glutathione metabolism, remained significant after multiple testing correction and were used for subsequent pathway and network analyses. (**E**) Enrichment map of GO biological processes. Each dot represents a GO term, colored based on the evidence from two individual (yellow: mRNA, blue: protein) and the combined (pink) analyses. The GO terms are clustered and labeled to reflect their shared biological theme.

Overall, there was a moderate Pearson correlation of r=0.31 (p<2.2e-16) between the log_2_FC at the mRNA and protein levels (**Figure 3B**). The directional p-value merging (DPM) method^22^ was used to integrate the p-values and directions of alcohol-induced changes in the RNA-seq and proteomics analyses, where genes in directional agreement (i.e., up-regulated or down-regulated in both analyses) were prioritized while those in directional conflict were penalized. The DPM method revealed 438 genes with a p-value <0.05 (**Figure 3C**, **Supplementary Table S4A**), among which 297 and 141 had the same and opposite fold-change directions, respectively, between the mRNA and protein levels. After multiple testing correction, 339 had an adjusted p-value <0.05 and were considered for subsequent analyses. **Figure 3D** shows the p-values and log_2_FC for the top penalized genes that were significant at both mRNA and protein levels, but penalized (merged p>0.05) due to directional conflict (**Supplementary Table S4B**). Although such directional conflicts may reflect post-transcriptional or post-translational regulation and warrant further mechanistic investigation, our integrated analysis here focused on concordantly regulated proteins. Accordingly, the “top enhanced proteins” panel in the same plot shows 8 proteins without a significant change in protein abundance but were identified as significant after combining the mRNA information with consistent FC direction (**Figure 3D**, **Supplementary Table S4B**). These genes/proteins, including Gsta2, Gsta4 and Gstp1 involved in glutathione metabolism, remained significant after multiple testing correction and were part of the subsequent pathway and network analyses.

The 339 significant genes identified with the DPM method were enriched in 168 GO biological processes (**Supplementary Table S4C**), where 134 were unique in this analysis (i.e., not identified with individual differential analysis of RNA-seq and proteomics data). The uniquely enriched biological processes were represented by several genes that were not significant in individual omics analyses (adjusted p>0.05) but prioritized due to their directional agreement between the mRNA and protein levels. These included Slc25a22 (mitochondrial glutamate carrier 1 involved in NADH metabolic and redox processes), Uqcrc1 (a complex III subunit required for aerobic electron transport and ATP synthesis-coupled electron transport, mitochondrial ATP synthesis coupled electron transport, etc.), and Ogdh (tricarboxylic acid (TCA) cycle enzyme linking acyl-CoA metabolism with NADH production and mitochondrial energy metabolism). Together, the enrichment of these mitochondrial genes highlights alterations in mitochondrial energy metabolism and redox homeostasis consistent with alcohol-induced mitochondrial dysfunction and oxidative stress^4^.

The enrichment map (**Figure 3E**, **Supplementary Table S4D**) facilitates a more systematic comparison of the biological processes identified with the RNA-seq, proteomics, and integrated analyses, where each node represents a GO:BP term, and an edge connects a pair of similar processes determined based on the overlapped genes. While both the RNA-seq and integrated analyses identified chronic alcohol-induced alterations in the multienzyme antioxidant defense system and retinoic and other vitamins, the integrated approach offered a more detailed characterization of ALD-related pathophysiology. Specifically, the integration revealed deeper perturbations in primary metabolic networks, including amino acid, lipid, nucleotide, and carbohydrate metabolism, as well as alcohol biotransformation and mitochondrial electron transport chain activity. On the other hand, several clustered biological processes that were uniquely identified with the RNA-seq analysis appeared to have limited relevance to ALD, including epithelial cell proliferation, cardiac muscle contraction, renal system development and response to metal ions. These results demonstrate that integrated multi-omics analysis provides a more robust signature of ALD than single-platform approaches. By filtering transcriptomic noise and identifying signals concordant with proteomic changes, this strategy prioritizes biologically and pathologically relevant pathways over transient fluctuations in gene expression.

### Protein network analysis

Besides the integrated mRNA/protein analysis, reanalysis of our previously published kinetic proteomics dataset^10^ revealed that alcohol significantly altered the stability of proteins involved in mitochondrial metabolism, energy production, and oxidative pathways, with coordinated changes observed across functional protein networks (**Supplementary Figure S3**, **Supplementary Table S5**), supporting alcohol-associated remodeling of mitochondrial and metabolic processes reflected at the levels of protein turnover and acetylome dynamics. To functionally characterize the differentially regulated proteome, we performed network-based integration and clustering analysis, identifying distinct functional modules enriched in oxidative stress signaling and glutathione metabolism. Proteins exhibiting altered expressions from integrated analysis or turnover rates were mapped onto a STRING protein-protein interaction (PPI) network, which was subsequently clustered to identify distinct functional modules. In the clustered network (**Supplementary Figure S4A**, **Supplementary Table S6A**), each node represents a protein, color-coded based on its mitochondrion score (0-5, higher values suggesting the more likely it is localized to mitochondrion), from the COMPARTMENTS database^23^ integrated in stringApp 2.0^24^. Of the seven major clusters identified (n ≥ 9 proteins), five (#2, #3, #4, #5 and #7) were highly represented by mitochondrial proteins (**Supplementary Figure S4A**). We next functionally interrogated these clusters using GO (Biological Process/Cellular Component), Reactome, and KEGG pathways, applying non-redundant significant terms for annotation in stringApp 2.0^24^ (**Figure 4A-B**, **Supplementary Figure S4B-H**, **Supplementary Table S6B-H**).

**Figure 4.**
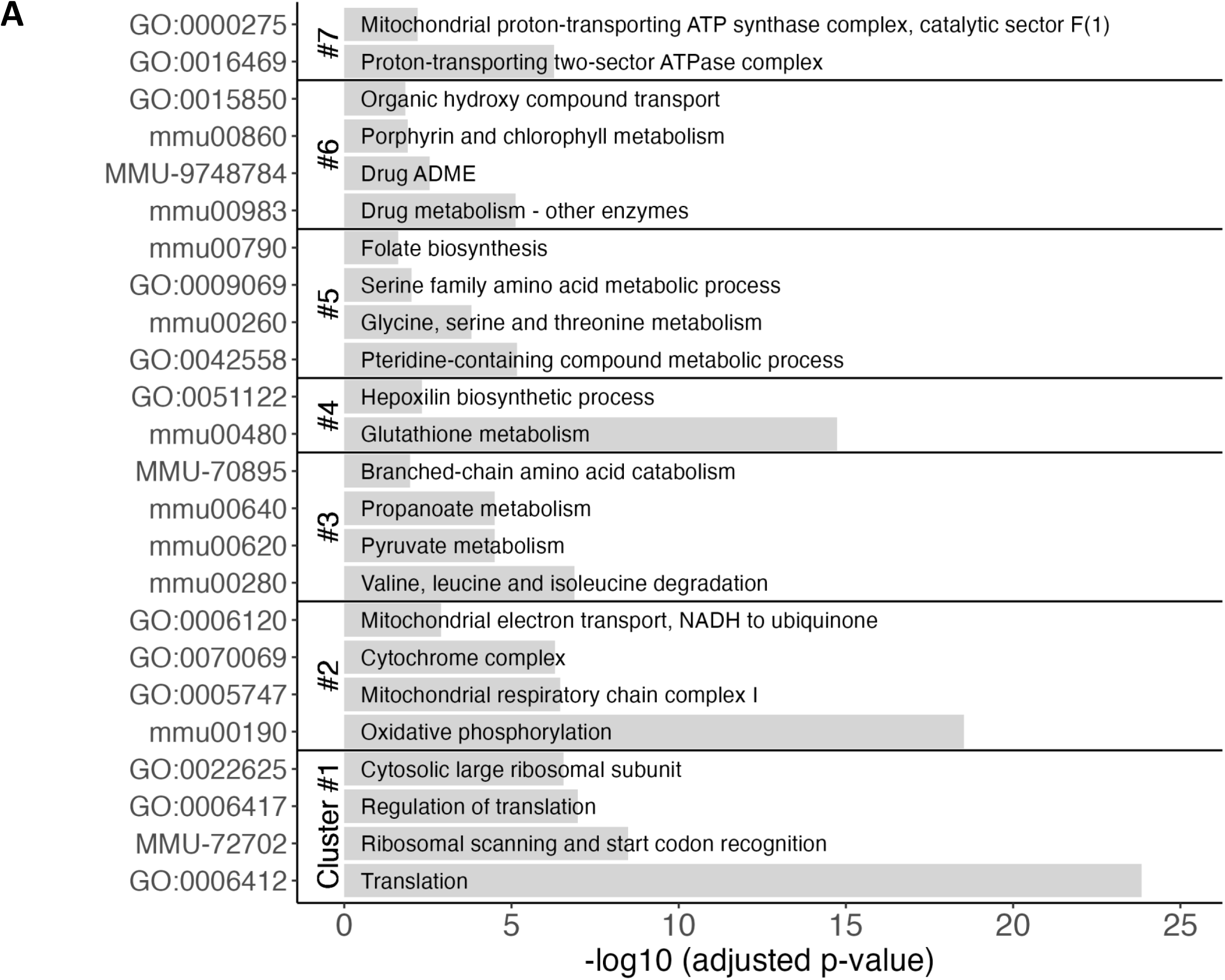
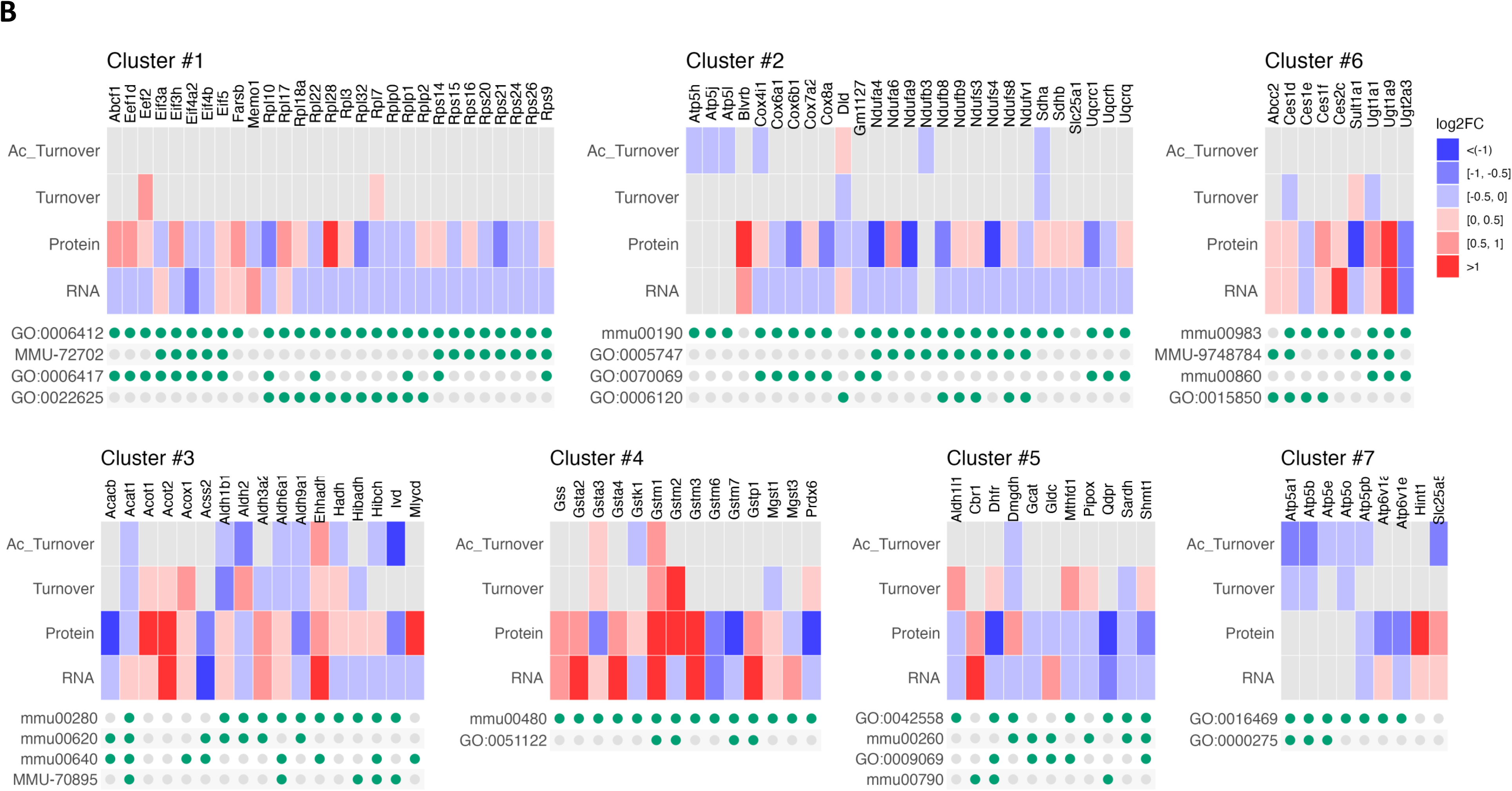
(**A**) Biological processes and molecular pathways associated with Clusters #1-7 in the interaction network of the proteins with altered expression or stability (**Supplementary Figure S4**). (**B**) Heatmap of the fold changes in mRNA, protein, turnover and acetylated turnover, for the proteins in each cluster, along with their association with the enriched biological processes and molecular pathways.

Cluster #1 (n=29 proteins) was overrepresented by ribosomal proteins and eukaryotic translation initiation factors, closely related to ribosomal scanning and start codon recognition and translation initiation (**Figure 4A-B**, **Supplementary Figure S4B**). Although turnover data were limited, both mRNA and protein levels were broadly decreased with alcohol exposure, suggesting compromised translational capacity consistent with cellular stress in alcohol-exposed liver^25^.

Cluster #2 (n=27) and Cluster #7 (n=9) were centered on mitochondrial proteins. Cluster #2 comprised essential subunits of Complex I (NADH dehydrogenase), Complex III (ubiquinol-cytochrome c reductase), and complex V (ATP synthase) (**Figure 4A-B**, **Supplementary Figure S4C**). Transcriptional downregulation of these genes, coupled with reduced turnover rates of acetylated proteins (Atp5h, Atp5j, Atp5l, Cox4i1, Ndufb3 and Sdha), indicates impaired mitochondrial electron transport. Similarly, Cluster #7 (n=9) included several ATP synthase subunits with decreased mRNA levels and attenuated turnover of both native and acetylated proteins (**Figure 4A-B**, **Supplementary Figure S4H**), collectively pointing to a deficit in mitochondrial bioenergetics in response to chronic alcohol exposure.

Metabolic impairment was further evidenced in Cluster #3 (n=17), which was enriched in several dehydrogenases associated with the metabolism of branched-chain amino acids (BCAAs), propionate, and pyruvate (**Figure 4A-B**, **Supplementary Figure S4D**). These changes, alongside the downregulation of genes involved in one-carbon metabolism and interconnected amino acid metabolism in Cluster #5 (n=11; **Figure 4A-B**, **Supplementary Figure S4F**), suggest coordinated disruption of mitochondrial respiratory function and methyl-donor metabolism. This pattern is consistent with alcohol-induced mitochondrial dysfunction and oxidative stress, which can perturb SAM-dependent methylation capacity and redirect sulfur amino acid metabolism toward antioxidant defense.

Conversely, we identified metabolic adaptive signatures in Clusters #4 and #6. The clustering of glutathione S-transferases (GSTs) in Cluster #4 (n=14) suggests a synchronized adaptive response to increased oxidative stress, as these enzymes play key roles in detoxifying reactive electrophiles and maintaining cellular redox homeostasis (**Figure 4A-B**, **Supplementary Figure S4E**). Furthermore, Cluster #6 (n=9) exhibited a coordinated upregulation of UDP-glucuronosyltransferases and carboxylesterases at the transcript and protein levels (**Figure 4A-B**, **Supplementary Figure S4G**), reflecting a robust hepatic response to enhance xenobiotic clearance and maintain metabolic homeostasis.

Together, these network-derived modules highlight a landscape of alcohol-induced altered one-carbon metabolism and mitochondrial failure countered by specific redox-active adaptations. This prompted us to investigate the transcriptional regulators underlying the proteomic changes and to determine whether these alterations contribute to metabolic reprogramming within the glutathione biosynthetic pathway through protein–metabolite interaction analysis.

### Transcription factor enrichment analysis

To identify potential upstream regulators of oxidative stress and glutathione pathways (Clusters #3-6), we used the ChEA3 platform to perform transcription factor (TF) enrichment analysis^26^ using the Literature ChIP-seq TF-target gene set library, prioritizing TFs whose target genes significantly overlapped with genes exhibiting altered expression and/or protein stability within these clusters (**Figure 4**).

TFs prioritized by ChEA3, including ESR1, EGR1, NFE2L2 (NRF2), GABPA, STAT5A, PPARG, SRF, NR0B1, HNF4α, and E2F1 (ranked 1-10), are broadly consistent with current understanding of ALD (**Figure 5A**, **Supplementary Table S7A**). The interaction network generated from ChEA3-integrated libraries (**Figure 5B**) revealed a highly interconnected transcriptional regulatory network centered on NRF2 (nuclear factor erythroid 2–related factor 2) as a major integration hub. With the exception of NR0B1 (nuclear receptor subfamily 0 group B member 1), which remained isolated, and STAT5A (signal transducer and activator of transcription 5A) and GABPA (GA-binding protein transcription factor subunit α), which were indirectly linked to NRF2 through EGR1-mediated stress-response signaling, all other TFs were connected to NRF2 through directed edges supported by ChIP-seq evidence curated within the ChEA3 platform, providing insight into potential upstream–downstream regulatory relationships within the network.

**Figure 5.**
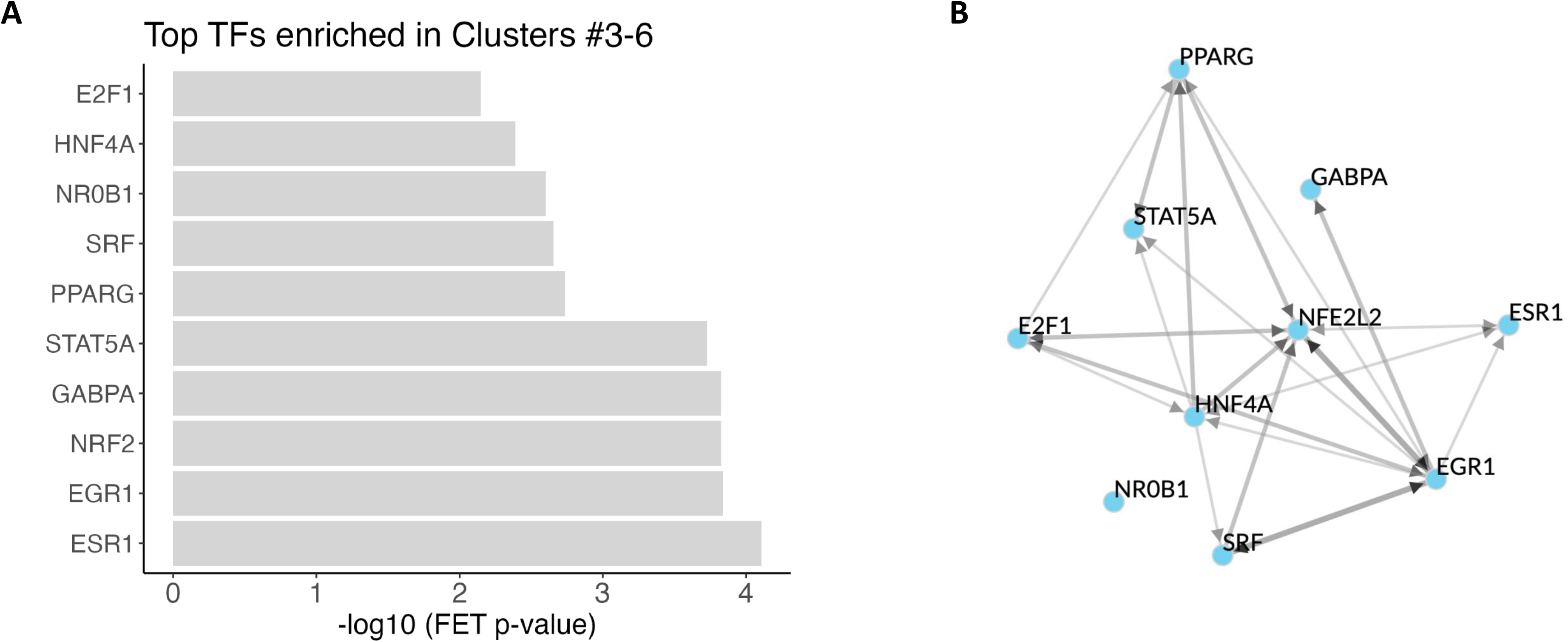
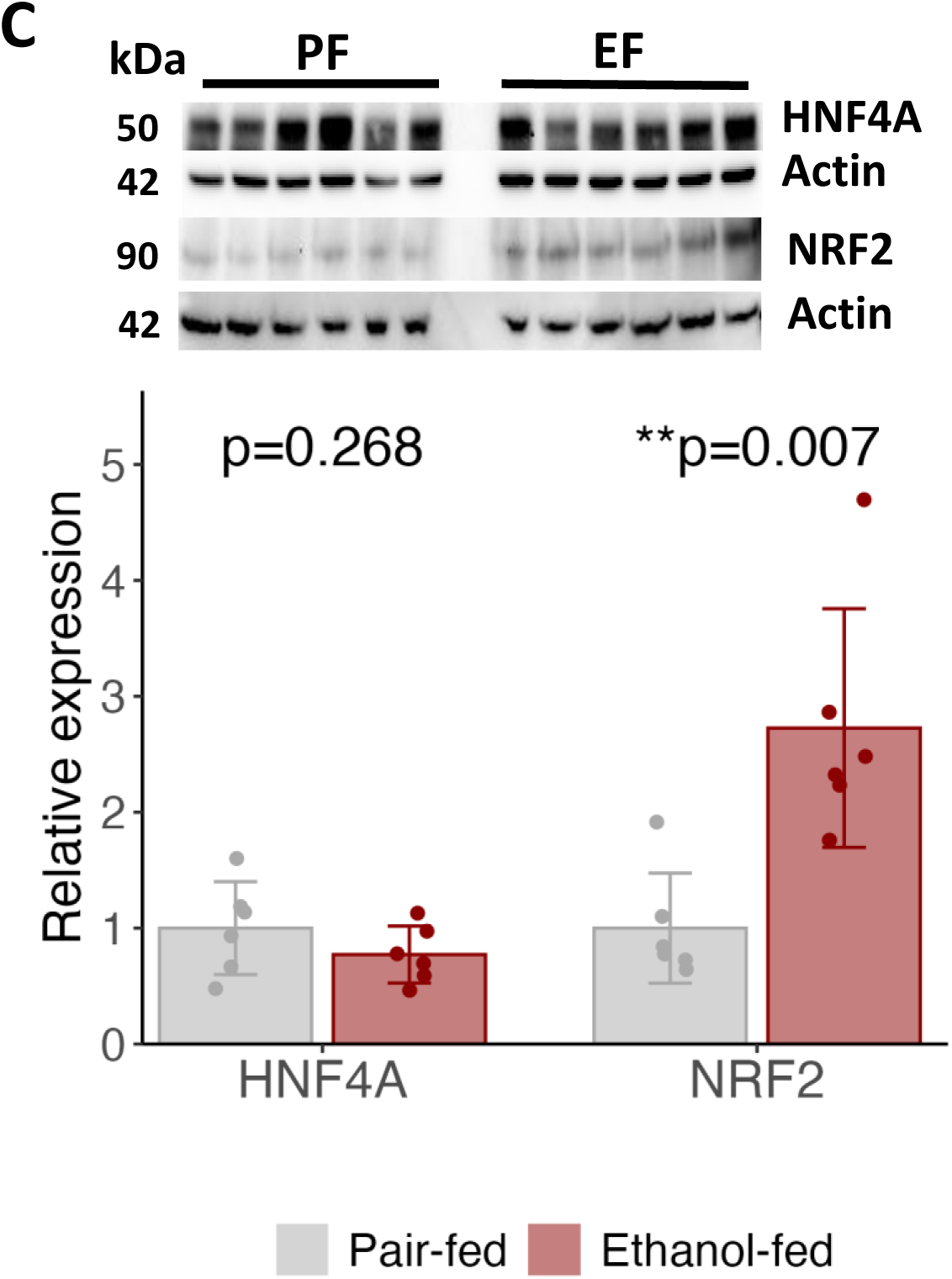
Transcription factors prioritized by ChEA3 for proteins with altered expression or turnover in Clusters #3-6 of the protein interaction network (Figure 4). (**A**) Top-ranked transcription factors (ranks 1-10). The x-axis represents the Fisher exact test p-values on a –log_10_ scale. (**B**) Interactions among the transcription factors shown in (**A**), based on evidence from ChEA3 libraries and, where applicable, supported by ChIP-seq data for directed interactions. (**C**) Western blot analysis of HNF4α and NRF2 levels in PF and EF mouse livers.

This network topology supports NRF2 as a central regulator coordinating the hepatic adaptive response to ethanol-induced oxidative stress and metabolic injury^27^. Within this network, EGR1 (early growth response 1), SRF (serum response factor), and E2F1 (E2F transcriptional factor 1) formed a stress-responsive module linking inflammatory activation, immediate-early gene signaling, and cell-cycle/repair pathways^28^. In parallel, PPARG (peroxisome proliferator-activated receptor γ), ESR1 (estrogen receptor α) and HNF4α converged toward NRF2-associated regulation of lipid metabolism, oxidative stress responses, and mitochondrial bioenergetics, processes profoundly disrupted during ALD progression^29^. STAT5A, which supports hepatocyte survival and metabolic homeostasis^30^ and GABRA, a key regulator of mitochondrial biogenesis and respiratory gene expression, further linked stress adaptation to metabolic remodeling within the network^31^.

A similar analysis of proteins with altered expression or turnover in Clusters #1–2 using the ChEA3 literature-derived ChIP-seq library identified c-Myc as a top enriched transcription factor (**Supplementary Figure S5A**, **Supplementary Table S7B**), highlighting its potential role as a central regulatory hub. This finding is biologically relevant because c-Myc has been implicated in HNF4α-mediated activation of NRF2-dependent cytoprotective pathways following acetaminophen-induced liver injury associated with glutathione depletion^32^. Moreover, impaired HNF4α signaling has been linked to hepatocellular injury, steatosis, and inflammation^18,33^, while reduced HNF4α activity is a hallmark of chronic liver disease progression^34^. HNF4α also plays a critical role in regulating sulfur amino acid metabolism and transsulfuration pathways required for glutathione synthesis^35^.

Given the central roles of HNF4α and NRF2 in chronic liver disease^33^ and anti-oxidant response^27^, respectively, we further examined their protein expression by immunoblot analysis (**Figure 5C**, **Supplementary Table S7C**). Consistent with clinical ALD studies showing progressive HNF4α loss^18^, chronic ethanol exposure in our murine model produced a trend toward reduced HNF4α protein expression, although the change was not statistically significant (**Figure 5C**). This likely reflects the relatively mild phenotype of our model, which represents early-stage ALD characterized by steatosis and modest inflammation^10^ rather than advanced hepatocellular dysfunction. In contrast, NRF2 protein expression increased following chronic ethanol exposure (**Figure 5C**), consistent with activation of antioxidant and glutathione-associated defense pathways in response to oxidative stress. These findings align with recent evidence demonstrating that HNF4α cooperates with c-Myc to enhance NRF2 activity and support antioxidant defense during oxidative stress^32^. Together, the convergence of the transcriptional interaction network on c-Myc and NRF2, combined with preserved HNF4α expression and NRF2 induction, suggests that the liver initially maintains hepatocyte metabolic identity while activating adaptive antioxidant and detoxification programs in response to chronic ethanol exposure.

### Integrated protein–metabolite analysis reveals an adaptive redox–metabolic axis

To determine whether these transcriptomics and proteomic alterations translate into functional adaptations within redox-regulating metabolic pathways, we constructed an integrated protein–metabolite interaction network centered on key metabolites from the methionine cycle and related pathways (S-adenosylmethionine, S-adenosylhomocysteine, homocysteine, methionine, 5-methyltetrahydrofolate, 5,10-methylenetetrahydrofolate, and tetrahydrofolate), transsulfuration (homocysteine, cystathionine, and cysteine), glutathione biosynthesis and utilization (gamma-glutamylcysteine, reduced glutathione, and oxidized glutathione), and ethanol metabolism (ethanol, acetaldehyde, acetate, NAD^+^, and NADH). These metabolites were queried against the STITCH database^36^ to find their interacting proteins, which were then filtered for specificity to liver tissues using the TISSUES resource^36^. Both STITCH search and filtering for tissue specificity were conducted in stringApp 2.0^24^. In the resulting network (**Supplementary Figure S6**), metabolites (squares) and their interacting proteins (circles), colored by changes in mRNA expression (left) and protein abundance (right) revealed highly connected hubs linking one-carbon and glutathione pathways. Upregulated proteins, including Gnmt, Chdh, and Dmgdh, formed central nodes connecting choline oxidation, methionine cycling, and methyl group flux.

This remodeling functionally converges on glutathione metabolism, encompassing coordinated regulation of GSH synthesis, utilization, and recycling (**Figure 6A**). Expression of key enzymes in de novo GSH synthesis is increased, including the rate-limiting catalytic subunit glutamate–cysteine ligase (Gclc, mRNA), its regulatory partner (Gclm, mRNA), and glutathione synthetase (Gss, mRNA and protein), supporting enhanced biosynthetic capacity. Concurrent upregulation of 5-oxoprolinase (Oplah, protein), which converts 5-oxoproline to glutamate within the γ-glutamyl cycle, further reinforces substrate availability for sustained GSH production. In parallel, elevated expression of glutathione peroxidases (Gpx3, Gpx4) and glutathione S-transferases (Gstp1, Gsta2, and Gsta4, mRNA and protein) is consistent with increased GSH utilization in antioxidant defense and xenobiotic detoxification. This is accompanied by upregulation of glutathione reductase (Gsr, mRNA and protein), indicating enhanced recycling of oxidized glutathione (GSSG) back to GSH, thereby maintaining redox buffering capacity. Notably, this coordinated anabolic and recycling program is coupled with suppression of GSH catabolic pathways, including reduced expression of γ-glutamyl cyclotransferase (Ggct, mRNA) and peptidases (Cndp2, Psma2, Psma5, and Psmb1, protein), limiting GSH degradation. Collectively, these changes define an integrated enzyme–metabolic network that sustains glutathione turnover and redox homeostasis, likely supported by increased flux through one-carbon metabolism and methionine utilization to preserve cysteine availability under conditions of oxidative stress.

**Figure 6.**
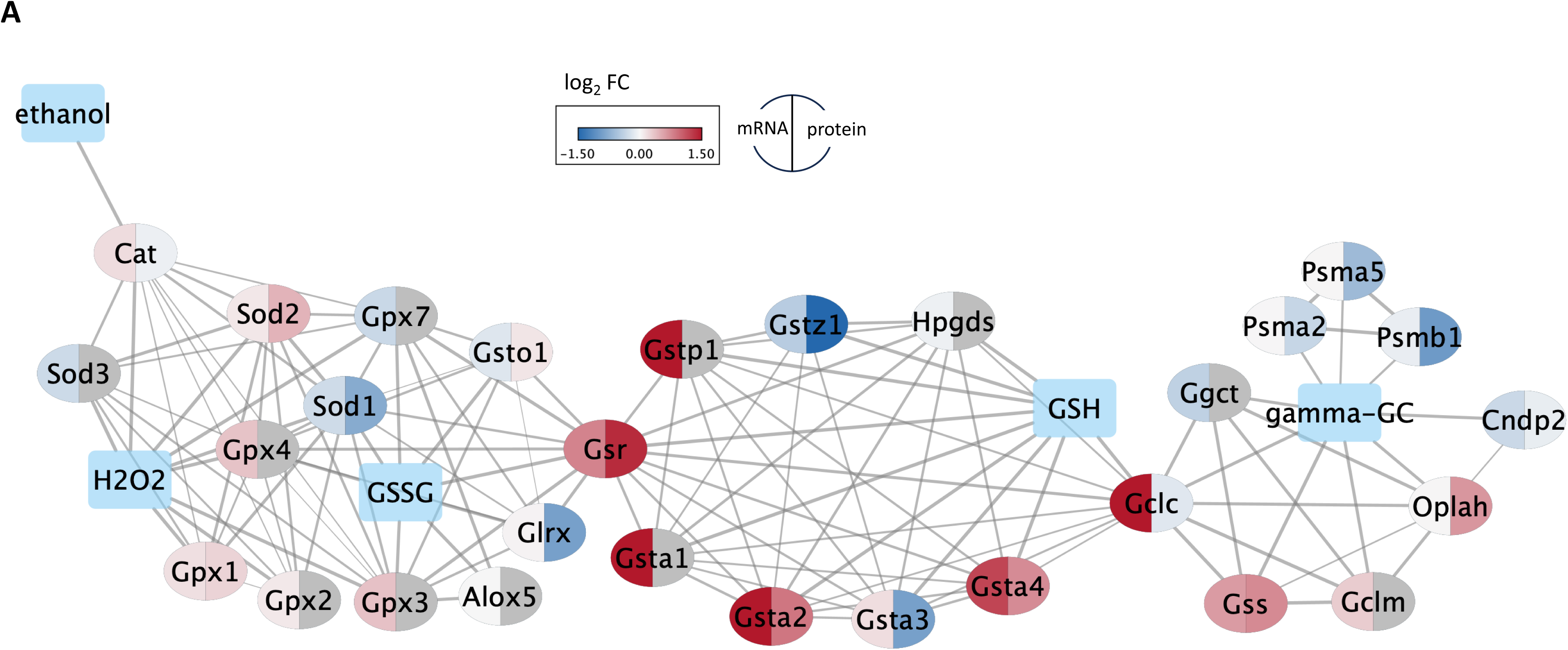
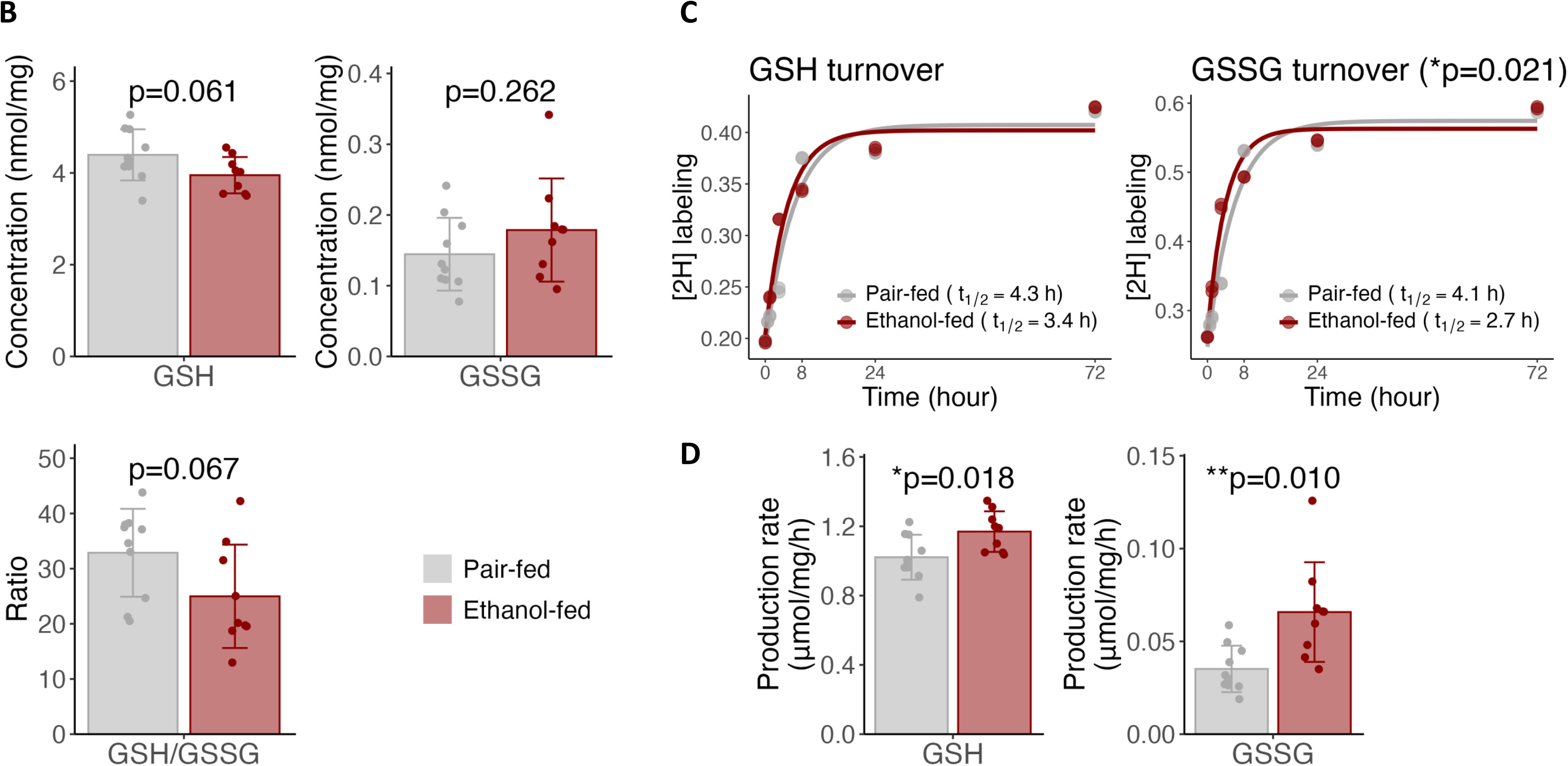
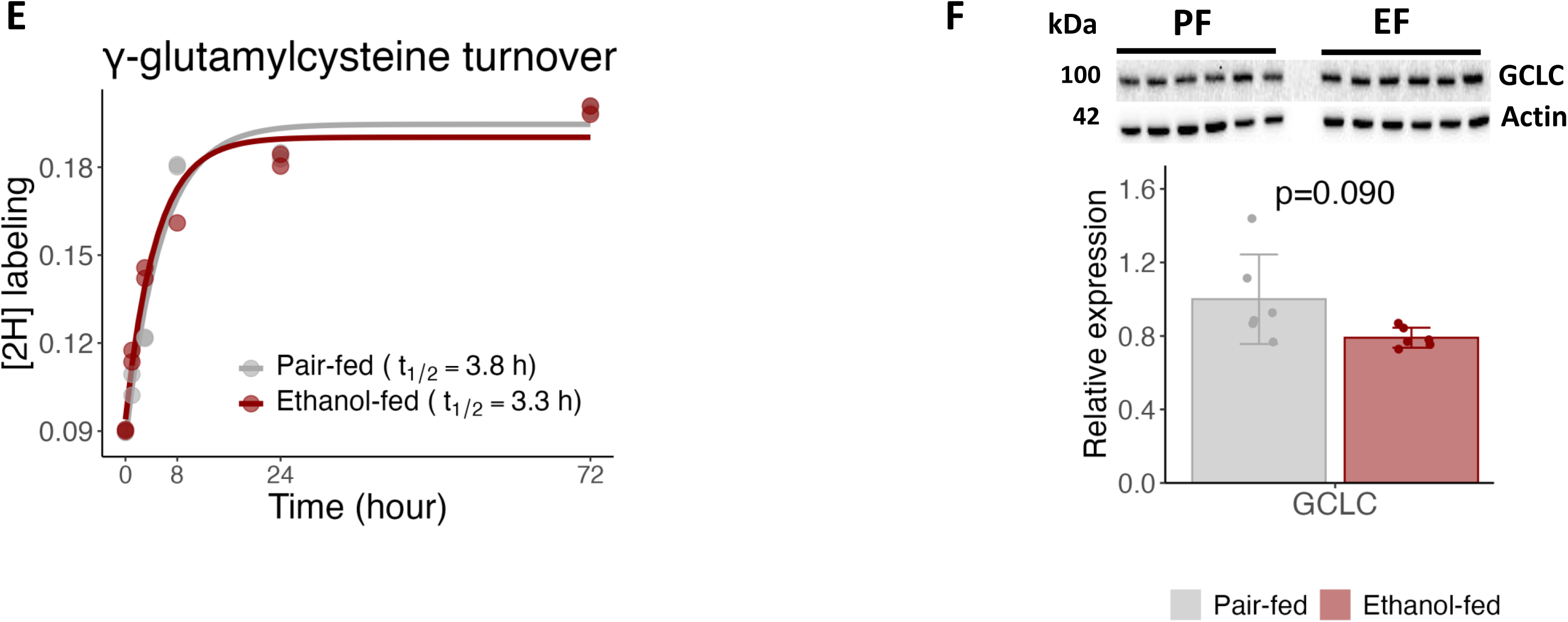
Chronic alcohol exposure enhances glutathione turnover and redox cycling. (**A**) Protein–metabolite interaction network comprising ethanol and hydrogen peroxide, an oxidative stress–related byproduct of ethanol metabolism, together with key metabolites of glutathione metabolism (squares), and their interacting proteins (circles), colored by changes in mRNA expression (left) and protein abundance (right). (**B-E**) Effect of chronic alcohol exposure on hepatic glutathione metabolism. (**B**) GSH, GSSG levels, and GSH/GSSH ratio in PF and EF mouse livers. (**C**) Chronic alcohol increased the turnover rate of GSSH, but not GSH. (**D**) Chronic alcohol increased the production rates of both GSSH and GSH. **(E)** Alcohol accelerates glutathione turnover without increasing γ-glutamylcysteine formation; ²H incorporation shows no enhancement of the GCL-catalyzed rate-limiting step in GSH synthesis. (**F**) Alcohol-enhanced GSH turnover is not significantly associated with changes in the GCL rate-limiting enzyme of GSH synthesis. Bar graphs and error bars in (**B**), (**D**), and (**F**) show sample mean ± SD, overlaid by individual data points. The means of PF and EF were compared using two-sample t-tests (*P<0.05, **P<0.01).

### Chronic alcohol exposure enhances glutathione turnover and redox cycling

Building on our previous observation that chronic alcohol exposure induces lipid peroxidation, as evidenced by increased hepatic malondialdehyde levels^10^, and guided by network-based analyses identifying oxidative stress responses and glutathione metabolism (**Figure 4**) as key affected modules, we next characterized hepatic glutathione redox dynamics using a ^2^H₂O metabolic labeling approach. As a universal tracer, ^2^H₂O enables simultaneous quantification of multiple biosynthetic pathways *in vivo*^37^. While extended labeling (up to 21 days) was used in our prior studies to determine protein and acetyl-protein turnover^10,13^, GSH and GSSG kinetics were assessed here using short-term labeling (≤3 days) to capture their rapid hepatic synthesis and turnover^38,39^. Quantification of reduced (GSH) and oxidized (GSSG) glutathione pools indicated a shift toward a more oxidized intracellular environment, reflected by a trend toward lower GSH levels (p=0.061) and a reduced GSH/GSSG ratio (p=0.067) (**Figure 6B**, **Supplementary Table S8A**).

Kinetic analyses further demonstrated that chronic alcohol exposure significantly accelerated glutathione flux. Alcohol increased the production rates of both GSH and GSSG and significantly elevated GSSG turnover (**Figure 6C-D**, **Supplementary Table S8B-C**). Although the fractional synthesis rate (FSR) of GSH exhibited only a nonsignificant upward trend (**Figure 6C**), alcohol exposure significantly elevated its production rates (**Figure 6D**), calculated by integrating FSR with the GSH pool size. The concurrent elevation of GSSG turnover suggests increased oxidation and recycling of glutathione, consistent with accelerated glutathione redox cycling and heightened antioxidant demand under alcohol-induced oxidative stress (**Figure 6D**).

GSH synthesis occurs via a two-step process. First, cysteine and glutamate are coupled to form γ-glutamylcysteine, catalyzed by glutamate–cysteine ligase (GCL). In the second step, glutathione synthetase (GS) catalyzes the addition of glycine to generate GSH. We, next, examined whether the alcohol-induced increase in glutathione synthesis was associated with changes in the first rate-limiting step of the pathway. Because γ-glutamylcysteine is a low-abundance and rapidly turning over intermediate in glutathione biosynthesis, its direct measurement in liver tissue is challenging. We therefore exploited fragment-ion–resolved isotope tracing, quantifying ²H incorporation into the γ-glutamylcysteinyl fragment of GSH by high-resolution MS/MS, which enables indirect but pathway-specific measurement of turnover at the GCL-catalyzed step *in vivo*. Notably, alcohol exposure did not significantly alter γ-glutamylcysteine turnover (**Figure 6E**, **Supplementary Table S8D**), indicating that the enhanced GSH flux is not driven by increased formation of this rate-limiting intermediate via GCL. Instead, these data suggest that alcohol promotes increased GSH utilization and recycling, consistent with elevated demand for peroxide detoxification and maintenance of redox homeostasis. In this context, stable γ-glutamylcysteine turnover suggests that precursor formation is preserved but not upregulated, whereas increased GSH flux is sustained downstream through enhanced GSSG-to-GSH recycling via GSR, γ-glutamyl cycle activity mediated by OPLAH, and increased consumption by GPX4 and glutathione S-transferases.

Consistent with these flux measurements and LFQ proteomics data, Western blot analysis showed no change in hepatic GCL protein abundance (**Figure 6F**, **Supplementary Table S8E**), further indicating that increased GSH turnover is driven primarily by enhanced utilization and recycling rather than upregulation of de novo synthesis. Supporting this interpretation, our prior metabolomics analysis in the same cohort demonstrated that chronic alcohol exposure increased hepatic levels of multiple amino acids, whereas glycine and cysteine, the key substrates for the rate-limiting step of GSH synthesis, remained unchanged, suggesting their preferential utilization to sustain GSH production^10^ (**Supplementary Figure S7A**). Together with the observed lack of changes in both GCL expression and γ-glutamylcysteine turnover, these findings indicate that alcohol-induced GSH dynamics are maintained through increased cycling and redox-dependent consumption rather than expansion of GCLC-dependent biosynthetic flux.

Consistent with enhanced NRF2 expression (**Figure 5C**) and its established role in redox homeostasis during oxidative stress^40^, these findings suggest that chronic alcohol exposure increases oxidative burden, leading to enhanced GSH utilization and oxidation. In response, compensatory antioxidant pathways are activated to maintain GSH homeostasis through sustained synthesis and increased reduction of oxidized glutathione (GSSG) back to GSH. However, these adaptive responses are insufficient to fully restore redox balance, as evidenced by the shift toward a more oxidized glutathione state.

### Alcohol-associated glutathione flux coincides with histone acetylation and methylation remodeling

Consistent with pronounced metabolic and redox remodeling, including enhanced glutathione flux and oxidative stress, alcohol exposure was associated with coordinated changes in histone PTMs that shift the chromatin landscape toward a more transcriptionally permissive state at stress-responsive loci. We observed site-specific increases in histone acetylation^10^, most notably at H3K18 and H3K23 (**Figure 7A**), known substrates of the p300/CBP acetyltransferase complex that are co-deposited at active promoters and enhancers^41^. The enrichment of H3K18ac and H3K23ac at antioxidant response element (ARE)-containing promoters is consistent with activation of NRF2-mediated stress-responsive transcriptional programs^42^.

**Figure 7.**
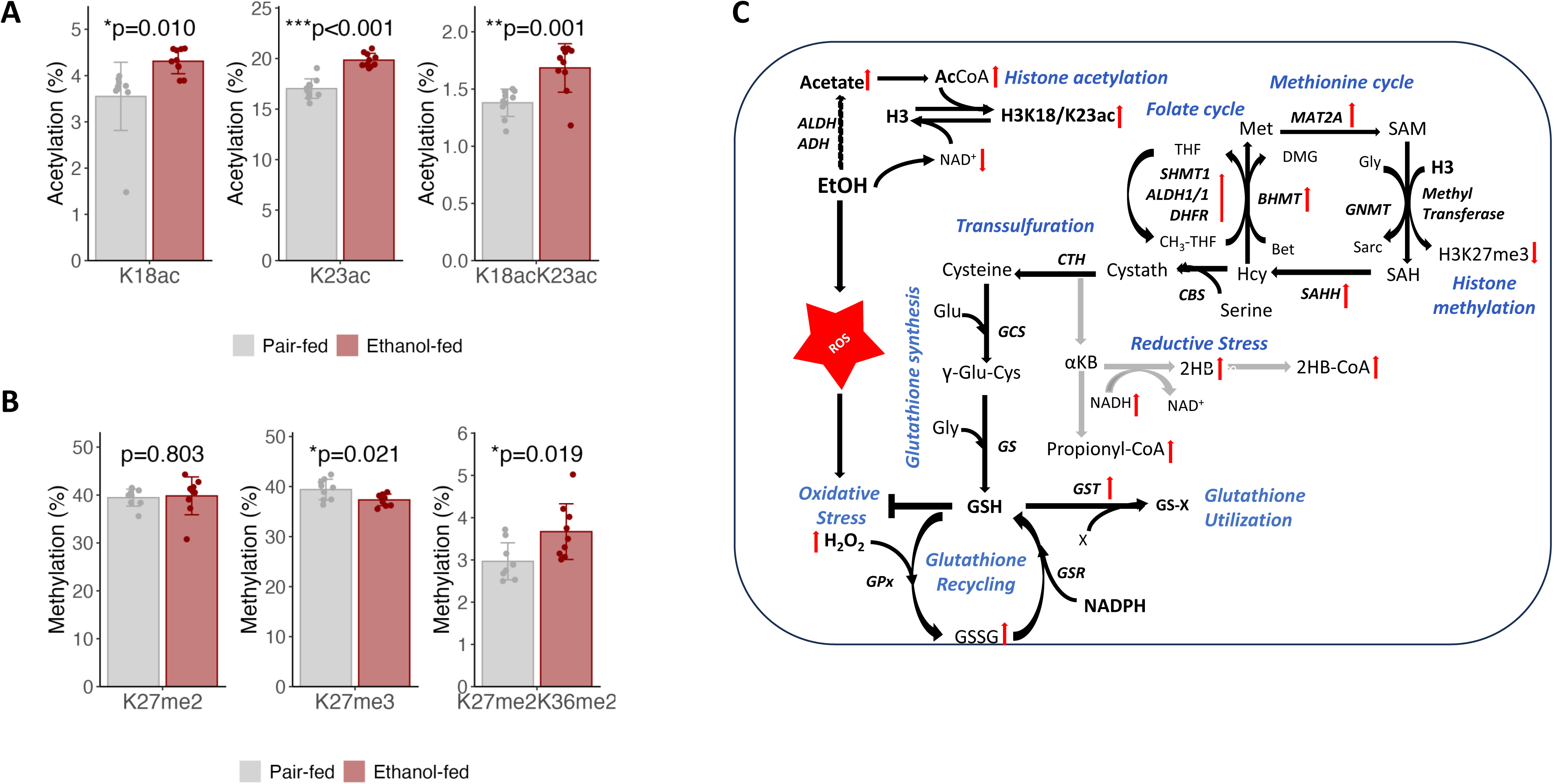
Alcohol-induced glutathione flux is linked to histone acetylation and methylation remodeling. Alcohol-driven redox and metabolic reprogramming, including increased glutathione (GSH) flux, is associated with coordinated changes in histone modifications, reflecting altered acetyl-CoA and SAM-dependent methylation capacity and a shift toward a more permissive chromatin state. **(A)** *Histone H3 acetylation.* Increased H3K18ac and H3K23ac, consistent with activation of stress-responsive transcription (e.g., NRF2 pathways). Data previously reported in Aghayev et al., 2025^10^; reanalyzed and visualized here. **(B)** *Histone H3 methylation.* Increased H3K36me2 with a decreased H3K27me3, indicating redistribution of methylation rather than global loss. **(C)** *Metabolic–epigenetic coupling.* Alcohol-induced metabolic reprogramming promotes a permissive chromatin landscape, primarily via enhanced histone acetylation driven by increased acetyl-CoA availability and reduced NAD⁺-dependent sirtuins. Concurrently, an adaptive antioxidant response marked by elevated glutathione (GSH) flux is accompanied by remodeling of histone H3 methylation, including a relative reduction in repressive H3 trimethylation marks. Gray arrows indicate increased flux through the transsulfuration pathway, and the consequent accumulation of associated side-products driven by elevated demand for cysteine and GSH synthesis. ALDH1: aldehyde dehydrogenase 1, BHMT: betaine–homocysteine methyltransferase, CBS: cystathionine beta-synthase, CTH: cystathionine gamma-lyase, DHFR: dihydrofolate reductase, DMG: dimethylglycine, GCS: γ-glutamylcysteine synthetase; E; electrophiles, GNMT: glycine N-methyltransferase; GS: glutathione synthase, GSH: reduced glutathione, GSSG: oxidized glutathione, GSR: glutathione reductase, GPx: glutathione peroxidase, GST: glutathione S-transferase, MAT2: methionine adenosyltransferase II, 2HB: 2hydroxybutyrate, 2HB-CoA: 2-hydroxybutyryl-CoA, SAH: S-adenosylhomocysteine, SAHH: S-adenosylhomocysteine hydrolase, SAM: S-adenosylmethionine, SHMT1: serine hydroxymethyltransferase 1, αKB: α-ketobutyrate.

In parallel, alcohol induced selective remodeling of histone methylation (**Figure 7B**, **Supplementary Table S9**). We observed an increase in the H3K27me2K36me2 peptide without a corresponding change in H3K27me2, indicating that the increase is driven by elevated H3K36me2 rather than altered H3K27 methylation per se. This was accompanied by a decrease in suppressive H3K27me3^43^ (**Figure 7B**), consistent with redistribution of methylation marks rather than global suppression. This pattern aligns with enhanced glutathione flux (**Figure 6**), characterized by increased GSH utilization supported primarily by recycling rather than expansion of GCL-dependent biosynthesis. This preferentially diverts homocysteine into the transsulfuration pathway to sustain cysteine supply and GSH turnover, potentially limiting remethylation to methionine and S-adenosylmethionine (SAM) generation. This shift is supported by increased transsulfuration-derived metabolites (e.g., 2-hydroxybutyrate (2HB), 2-hydroxybutyryl-CoA (2HB-CoA), and propionyl-CoA) (**Supplementary Figure S7B**).

Reduced SAM availability is known to preferentially constrain higher-order methylation^44^, reflected here by decreased H3K27me3 and a relative enrichment of dimethylated species, consistent with a shift toward a more permissive chromatin state (**Figure 7C**)^45^. Collectively, these findings demonstrate that alcohol-induced metabolic and redox remodeling drives coordinated histone acetylation (increased H3K18ac and H3K23ac) and redistribution of methylation marks (decreased H3K27me3 and increased H3K27me2K36me2), promoting NRF2-mediated stress-responsive gene expression.

## DISCUSSION

Hepatic ethanol metabolism generates ROS primarily via the microsomal ethanol-oxidizing system, particularly CYP2E1, with a minor contribution from peroxisomal catalase–mediated oxidation^46,47^. Acetaldehyde, produced by these pathways and mitochondrial alcohol dehydrogenase (ADH), further exacerbates oxidative stress by impairing the ETC and promoting mitochondrial dysfunction. Our previous work showed that this dysfunction is associated with increased protein acetylation, selectively impairing turnover of mitochondrial, but not cytosolic, proteins^10^. This metabolic stress drives lipid peroxidation and results in mild hepatocellular injury in the absence of significant inflammation or fibrosis. Here, to define mechanisms of hepatic resilience, we applied transcriptomic and integrated multi-omics approaches and identified mitochondrial oxidative phosphorylation as a primary target in early-stage ALD. Concurrently, we identified the anti-oxidant glutathione system as a key compensatory pathway, characterized by sustained biosynthesis alongside enhanced utilization and recycling. Metabolic flux analysis and molecular profiling demonstrate that chronic ethanol exposure induces a high-flux glutathione cycle, characterized by increased glutathione turnover and recycling. This adaptive remodeling of redox metabolism precedes overt liver injury and likely serves to buffer oxidative stress, thereby limiting disease progression.

Our transcription factor enrichment analyses identified NRF2 as a putative upstream regulator of the observed anti-oxidant stress response, and several other TFs including ESR1 and HNF4α involved in sexual dimorphism, metabolism, inflammatory signaling. The enrichment of ESR1 as an upstream regulator suggests a female-specific pathway contributing to alcohol-induced hepatic inflammation and metabolic dysregulation in the female mice used in this study, distinct from estrogen receptor signaling profiles reported in male mice^48^. Given the known role of ERα in hepatic lipid metabolism, and redox regulation, ESR1 enrichment may indicate that estrogen-responsive transcriptional programs contribute to the adaptive and/or maladaptive hepatic response to chronic ethanol exposure. This is particularly relevant in female mice, where estrogen signaling has been implicated in the sexual dimorphism of ALD and may intersect with NRF2-dependent antioxidant pathway^49^.

As a master regulator of hepatic metabolic homeostasis, HNF4α controls gene programs governing gluconeogenesis, bile acid synthesis, and lipid, cholesterol, amino acid, and sulfur metabolism, and is highly sensitive to redox-dependent modulation^16, 35,50^. Its progressive functional decline is a hallmark of chronic liver diseases, including metabolic dysfunction–associated steatotic liver disease/steatohepatitis (MASLD/MASH), hepatocellular carcinoma, and alcoholic hepatitis^34^. ROS generated during ethanol metabolism can promote PKC-mediated phosphorylation of HNF4α, leading to its cytoplasmic sequestration and enhanced proteasomal degradation^51^. Our acetyl-proteomic and turnover analyses further implicate mitochondrial dysfunction as a proximal driver of oxidative stress. Alcohol exposure induced widespread protein hyperacetylation accompanied by a selective reduction in mitochondrial protein turnover, whereas cytosolic turnover was largely preserved^10^. This pattern is consistent with lysine hyperacetylation disrupting mitochondrial proteostasis, impairing respiratory chain function, and compromising redox balance. The resulting mitochondrial stress is expected to elevate ROS, thereby reinforcing HNF4α destabilization while concurrently stabilizing and activating NRF2-dependent antioxidant programs.

Mechanistically, oxidative stress activates NRF2 through disruption of the KEAP1–NRF2 repressor complex^52^. Under basal conditions, NRF2 is sequestered by Keap1 and targeted for proteasomal degradation; oxidative stress disrupts this interaction, promoting NRF2 stabilization, nuclear translocation, and activation of antioxidant response element (ARE)-driven genes^40^. Accordingly, our data indicate that early ethanol exposure suppresses HNF4α-dependent metabolic regulation while activating NRF2-driven antioxidant programs, reflecting a redox-dependent shift in transcriptional control. Under basal conditions, HNF4α supports expression of genes involved in sulfur amino acid metabolism and glutathione biosynthesis^35^; however, oxidative stress preferentially engages NRF2-dependent pathways that enhance cellular antioxidant capacity^53^. Consistent with these mechanisms, we observed increased expression and turnover of key glutathione pathway components, including Gclc *and* Gss, the enzymes in GSH synthesis, and Gsr, which regenerates GSH from GSSG, and GSH-utilizing detoxification enzymes such as Gsta2 and Gstp1, many of which are established NRF2 targets, along with increased NRF2 expression.

Isotope tracer–based flux analyses demonstrate that these transcriptional and proteomic adaptations are accompanied by markedly accelerated turnover of both reduced (GSH) and oxidized (GSSG) glutathione. High-resolution mass spectrometry further indicates that enhanced GSH turnover is maintained without expansion of the GCL-dependent, cysteine-limiting step, alcohol exposure primarily enhances GSH utilization, which is counterbalanced by increased recycling through GSSG reduction. Thus, regulation of GSH flux occurs predominantly at the levels of utilization and recycling rather than de novo synthesis. Collectively, these findings indicate that the hepatic antioxidant system operates in a high-flux, nonequilibrium state, where sustained GSH synthesis and enhanced recycling compensate for increased consumption, thereby maintaining redox homeostasis during the early stages of ALD.

Multiple orthogonal datasets support the existence of this adaptive high-flux glutathione cycle. First, transcriptomic profiling reveals coordinated upregulation of genes involved in GSH biosynthesis and recycling (Gclc, Gss, Oplah, Gsr), consistent with increased synthetic and reductive capacity. Second, metabolomic analyses show that glycine and cysteine, the principal substrates for GSH synthesis, remain stable despite broader alterations in amino acid pools, suggesting preferential channeling of these substrates into glutathione production. Third, induction of phase II detoxification enzymes (Gsta2, Gstp1) indicates increased demand for GSH-dependent conjugation reactions to neutralize lipid peroxidation–derived electrophiles. Together, these findings define a coordinated, flux-driven adaptation linking mitochondrial dysfunction, redox signaling, and glutathione metabolism during early alcohol-induced metabolic stress.

Our results also show that alcohol-induced oxidative stress and the associated increase in glutathione flux are associated with epigenetic remodeling. Ethanol metabolism disrupts one-carbon metabolism by promoting oxidative stress–driven diversion of homocysteine into the transsulfuration pathway to sustain GSH synthesis^54^. This shift limits homocysteine remethylation to methionine, thereby impairing methionine cycling and reducing SAM availability^54^. Chronic ethanol exposure is also known to induce site-specific alterations in histone methylation, including H3K4 and H3K9 marks, reflecting disruption of SAM-dependent methylation capacity^55^. In parallel, ethanol metabolism increases hepatic acetyl-CoA availability and increases NADH at the expense of NAD⁺, which inhibits NAD⁺-dependent deacetylases (e.g., sirtuins), thereby promoting global and site-specific protein and histone hyperacetylation, a well-established feature of ALD^10,16^. Our data extend these observations by demonstrating a coordinated shift toward a more transcriptionally permissive chromatin landscape. This shift is characterized by increased histone H3K18 and H3K23 acetylation, reduced H3K27me3, a canonical Polycomb-associated repressive mark^56^, and a concomitant increase in H3K36me2, inferred indirectly from the increased H3K27me2K36me2 signal in the absence of changes in H3K27me2. Together, these changes are consistent with redistribution of histone methylation states rather than global suppression and align with concurrent metabolic shifts: enhanced acetyl-CoA availability favoring lysine acetylation, coupled with constrained SAM-dependent methylation capacity limiting higher-order methylation. Although global methylation was not directly measured, the reciprocal changes in acetylation and methylation marks support metabolically driven epigenomic reprogramming under alcohol-induced stress. The alcohol-induced epigenetic changes support a coordinated cellular response to redox–metabolic stress. Increased H3K18ac and H3K23ac are consistent with p300/CBP–NRF2 activation at ARE-containing loci^57^, decreased H3K27me3 and relative enrichment of H3K36me2, indicative of Polycomb derepressing^58^. These complementary remodeling layers, one driven by elevated acetyl-CoA and impaired sirtuin activity, the other by transsulfuration-linked methylation remodeling^54^, may act in parallel to reprogram transcription toward antioxidant defense and metabolic adaptation, while potentially predisposing to maladaptive gene expression relevant to alcohol-associated disease progression.

Notably, this shift toward a more permissive chromatin state was accompanied by robust induction of NRF2, a central regulator of antioxidant and cytoprotective responses, whereas HNF4α exhibited only a modest, non-significant decrease. These findings suggest that alcohol-induced disruption of hepatic metabolic identity is driven primarily by selective activation of redox-responsive transcriptional programs rather than broad suppression of hepatocyte gene expression. In particular, NRF2 activation appears to coincide with chromatin remodeling characterized by increased activating histone acetylation and redistribution of methylation marks, potentially facilitating induction of antioxidant defense pathways during early alcohol exposure. Although metabolic homeostatic programs may become partially attenuated during this transition, the overall response is dominated by NRF2-mediated adaptive stress signaling. Functionally, this coordinated epigenetic and transcriptional remodeling likely represents an early compensatory mechanism that enhances redox buffering capacity; however, persistent remodeling may ultimately reinforce transcriptional programs associated with metabolic dysfunction and liver injury.

In summary, our integrated multi-omics analysis demonstrates that chronic alcohol exposure induces early hepatic metabolic and epigenetic remodeling prior to overt liver injury. This pre-pathological state is marked by a high-flux glutathione cycle, in which redox balance is maintained through increased turnover. This adaptation is accompanied by coordinated metabolic–epigenetic interactions, where increased acetyl-CoA availability and limited methyl donor capacity, favor site-specific histone acetylation and redistribution of methylation marks^52,59^. Collectively, these findings support activation of NRF2-mediated antioxidant and stress-response pathways during early alcohol exposure, promoting metabolic adaptation and redox defense^32^. Although modest attenuation of HNF4α signaling may contribute to selective changes in hepatic metabolic programs, the mechanisms linking NRF2-dependent redox-sensitive signaling, chromatin remodeling, and progressive metabolic dysfunction remain to be defined.

## METHODS

### Materials

Common solvents and chemicals, including GSH, GSSG, dithiothreitol (DTT), N-ethylmaleimide (NEM) were purchased from Sigma-Aldrich (St. Louis, MO).

### Experimental design and mouse model

#### Chronic alcohol exposure and ^2^H_2_O-metabolic labeling

To quantify proteome and acetylome dynamics, mice were administered ²H₂O via liquid diets over time courses extending up to 21 days, and liver samples were collected at defined intervals following initiation of labeling, as previously described^10^. For assessment of glutathione kinetics, liver samples harvested at 0, 1, 2, 4, 8, 24, and 72 hours of ²H₂O metabolic labeling were analyzed. Body water enrichment was achieved by an initial ²H₂O–saline bolus followed by provision of 6% ²H₂O in drinking water, resulting in steady-state enrichments of 3.5 ± 0.6% and 3.5 ± 0.5% in the pair-fed (PF) and ethanol-fed (EF) groups, respectively. Animal experiments used in this study were described previously. Briefly, female C57BL/6 mice were obtained from The Jackson Laboratory (Bar Harbor, ME). All procedures were approved by the Institutional Animal Care and Use Committee. To evaluate the effects of chronic alcohol exposure, 2-month-old mice were pair-fed a modified Lieber–DeCarli liquid diet containing ethanol or an isocaloric maltose dextrin control for 4 weeks using a stepwise feeding protocol as previously described. Ethanol concentration was initially 1% (wt/vol; 7% of total calories) and gradually increased to 6% (36% of total calories). Control mice were pair-fed based on the daily intake of ethanol-fed mice (average intake ∼10 g·day⁻¹·mouse⁻¹). To quantify proteome and acetylome dynamics, mice were administered ²H₂O via liquid diets over time courses extending up to 21 days, as previously described. For assessment of glutathione kinetics, liver samples were collected at 0, 1, 2, 4, 8, 24, and 72 hours following initiation of ²H₂O labeling. Body water enrichment was achieved by an initial ²H₂O–saline bolus followed by provision of 6% ²H₂O in drinking water, resulting in steady-state enrichments of 3.5 ± 0.6% and 3.5 ± 0.5% in the pair-fed (PF) and ethanol-fed (EF) groups, respectively. At the end of the feeding period, mice were anesthetized and liver tissues were harvested. Whole-tissue lysates were prepared for protein and RNA analyses.

### Analytical methods

#### Total body water enrichment

The [^2^H]-enrichment of total body water was measured using a modified acetone exchange method^60,61^. Briefly, 5 μL of plasma, run in parallel with calibration standards, was incubated with 5 μL of 10 N KOH and 5 μL of acetone in a 2 mL sealed GC vial at room temperature for 4 h. Headspace acetone (1 μL) was analyzed by GC–MS (Agilent) in electron impact mode, monitoring ions m/z 58 (M0), 59 (M1), and 60 (M2). ²H enrichment was calculated from the calibration curve regression.

#### Sample preparation for hepatic glutathione analysis

Reduced and oxidized glutathione were quantified simultaneously by LC–MS following derivatization of GSH with N-ethylmaleimide (NEM). A seven-point calibration curve (0–100 nM) was generated using authentic GSH standards, each spiked with 20 nM isotopically labeled internal standard ([¹³C₂, ¹⁵N]-glycine, M+3 GSH).

Briefly, ∼20 mg of liver tissue was homogenized in 400 µL of 10 mM NEM in methanol containing the internal standard (200 nM in 20 µL). Chloroform (400 µL) was added, and samples were homogenized for 30 s, followed by addition of 400 µL of 10 mM NEM in water and further homogenization for 40 s to ensure complete extraction. Samples were incubated at room temperature for 15 min and centrifuged for 20 min. The aqueous phase was collected, dried under nitrogen, reconstituted in 80 µL water, and subjected to LC–MS analysis.

#### Glutathione assay by mass-spectrometry

Glutathione was analyzed using a Vanquish UHPLC system (Thermo Fisher Scientific, CA) coupled to a Q Exactive™ Plus Hybrid Quadrupole-Orbitrap™ mass spectrometer (Thermo Fisher Scientific, CA). Five μL of sample was injected onto a C18 reversed-phase column (75 μm × 15 cm, 2 μm, 100 Å; Thermo Fisher). Separation was performed at 0.30 mL/min using mobile phase A (ACN with 0.2% formic acid) and mobile phase B (60% acetonitrile, 40% isopropanol, 0.2% formic acid). After 2 min equilibration at 0% B, a linear gradient to 50% B over 10 min was applied, followed by an increase to 90% B in 1 min and a 5 min hold at 90% B.

Mass spectrometry analysis of derivatized GSH (GSH-NEM) and GSSG was performed in data-dependent acquisition (DDA) mode with a full profile MS scans at 70,000 resolution (200 *m/z*) between 55 and 855 *m/z*. MS/MS spectra were collected in data-dependent acquisition mode for the 3 most abundant precursor ions with an isolation window of 3 *m/z* at 17,500 resolution (at 200 *m/z*) with mass range starting at *m/z* 90. Higher-energy collisional dissociation (HCD) was performed at a normalized collision energy of 25%. The precursor ion masses were dynamically excluded from MS/MS analyses for a duration of 10 sec. MS and MS/MS spectra were acquired for inject time of 100 msec with the automatic gain control (AGC) target set at 3.0 x 10^6^ and 8.0 x 10^3^ ions for MS and MS/MS scans, respectively. Injection time for MS/MS scan does not exceed 50 msec. This DDA approach enabled acquisition of ∼50–80 MS1 (precursor) survey scans.

²H enrichment of the γ-glutamylcysteinyl fragment of GS-NEM was quantified using parallel reaction monitoring (PRM) on a Q Exactive Plus (Thermo Fisher Scientific). PRM was selected because it provides high accuracy and reproducibility for isotope-resolved fragment ion analysis^62^. To enhance sensitivity, collision energy was optimized to enhance detection of labeled fragment ions. Targeted GS-NEM precursor ions (*m/z* 433.1388) were isolated using a 3 *m/z* window with 0.6 *m/z* offset and fragmented by HCD at 35% normalized collision energy. MS/MS spectra were acquired at a resolution of 35,000, with an AGC target of 1×10⁶ and a maximum injection time of 200 ms. Under these conditions, approximately 100–150 MS/MS scans per acquisition cycle were obtained for the γ-glutamylcysteinyl fragment ion (*m/z* 358.1061), enabling robust quantification of its mass isotopomer distribution and precise assessment of ²H incorporation.

#### Glutathione turnover analysis

Time-course ^2^H enrichment of GSH and GSSG, and their fragment ions was used to determine their turnover rate constant. The fractional synthesis rate (FSR) was estimated using a single-compartment model by fitting the time course of total labeling *E*(*t*) to an exponential rise equation:

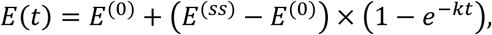

where *E^(0)^* and *E^(ss)^* are the baseline and plateau enrichments, respectively, and *k* represents the FSR.

Total ^2^H-labeling (*E(t)*) was calculated as the sum of molar percent enrichment (MPE) of the measured individual isotopomers (M₀–M_2_ for molecular ions and M₀–M_1_ for fragment ions). MPE for each isotopomer was determined from peak areas:

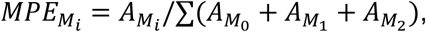

where *A*_*M*_ is the area of the *i*^*t*ℎ^ isotopomer.

Glutathione production rate (PR) was calculated as:

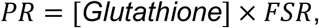

and expressed as µmol·gww⁻¹·h⁻¹, where glutathione concentration is given as µmol per g wet liver weight (gww).

#### Proteomics and acetylomics analysis

Proteomics data, including label-free quantification (LFQ) of protein abundance and global as well as acetylated protein turnover, used for integrative analyses were generated as previously described^10^. Briefly, liver samples were analyzed by high-resolution LC–MS/MS on a Q Exactive Orbitrap platform, and proteins were quantified using LFQ. Protein and site-specific acetylation turnover rates were determined using ^2^H₂O metabolic labeling, enabling simultaneous assessment of protein abundance, turnover, and acetylation dynamics. For LFQ, raw data were processed using established pipelines (e.g., MaxQuant) with a 1% false discovery rate (FDR) at the peptide and protein levels. Protein and acetylated protein turnover rates were quantified from time-course ^2^H labeling of native and acetylated peptides using a single-compartment kinetic model. Detailed experimental procedures, including sample preparation, acquisition parameters, and turnover calculations, are described in the original publication^10^.

#### Metabolic profiling

Targeted metabolic profiling was performed to quantify amino acids, acyl-CoAs, acyl-carnitines, and 2-hydroxybutyrate^10^. Amino acids and 2-hydroxybutyrate were analyzed by gas chromatography–mass spectrometry (GC–MS)^63^, while acyl-CoAs and acyl-carnitines were quantified using LC–MS/MS on a SCIEX platform^64,65^. Metabolites were extracted from liver tissue using established protocols and quantified using appropriate internal standards and calibration curves. Data were normalized to tissue weight and used for downstream integrative analyses with transcriptomic and proteomic datasets.

#### Total RNA extraction and sequencing

Total RNA was extracted from mouse liver tissue using the Monarch® Total RNA Miniprep Kit (NEB, T2010S) according to the manufacturer’s instructions. Briefly, ∼20 mg of liver tissue was homogenized in lysis buffer, and the lysate was clarified by centrifugation. RNA was bound to a silica membrane, followed by on-column DNase I treatment to remove genomic DNA contamination. After sequential wash steps, RNA was eluted in nuclease-free water. RNA concentration was determined by absorbance at 260 nm (A260), and purity was assessed using the A260/A280 ratio measured on a Synergy H1 Hybrid Reader (BioTek, Winooski, VT). Only samples with A260/A280 ratios between 1.9 and 2.1 were used for downstream analysis.

RNA sequencing was performed by LC Sciences (Houston, TX, USA). RNA integrity was further assessed, and only high-quality samples with RNA Integrity Number (RIN) > 7.0 were used for library preparation. Poly(A)+ mRNA was isolated from 5 µg of total RNA using Dynabeads Oligo(dT) (Thermo Fisher Scientific) with two rounds of purification. Purified mRNA was fragmented using divalent cations at elevated temperature (94°C for 5–7 min; NEB Magnesium RNA Fragmentation Module). First-strand cDNA was synthesized using SuperScript™ II Reverse Transcriptase (Invitrogen), followed by second-strand synthesis using *E. coli* DNA polymerase I, RNase H (NEB), and dUTP incorporation to generate strand-specific libraries. The resulting double-stranded cDNA was end-repaired, A-tailed, and ligated to dual-index adapters containing T-overhangs. Libraries were size-selected using AMPure XP beads. Uracil-DNA glycosylase (UDG; NEB) treatment was applied to degrade the dUTP-containing second strand, preserving strand specificity. Libraries were PCR-amplified (8 cycles: 98°C for 15 s, 60°C for 15 s, 72°C for 30 s; initial denaturation at 95°C for 3 min and final extension at 72°C for 5 min). The final libraries had an average insert size of 300 ± 50 bp. Paired-end sequencing (2 × 150 bp) was performed on an Illumina NovaSeq™ 6000 platform following the manufacturer’s protocols.

#### Immunoblot analysis

Liver tissue lysates were prepared and ∼30 µg of protein was separated by SDS–PAGE and transferred to polyvinylidene fluoride membranes. Membranes were blocked with 5% nonfat milk in TBS-T and incubated overnight at 4°C with rabbit polyclonal antibodies against GCL (Cell Signaling Technology, #48005), NRF2 (Abcam, #ab325240), and HNF4α (Invitrogen, MA1-199). After washing, membranes were incubated with HRP-conjugated donkey anti-rabbit IgG (GE Healthcare). Protein bands were detected using enhanced chemiluminescence (GE Healthcare) and quantified by densitometry. Protein abundance was normalized to the corresponding loading control.

### Quantification and statistical analysis

#### Differential gene expression analysis

Differential expression (DE) analysis for the RNA-seq dataset was performed using DESeq2 V1.48.1^66^. A pre-filtering was performed to keep only genes (n=17,003) that have a count of at least 10 for at least 6 samples. The adaptive prior shrinkage estimator from the apeglm package^67^ was used to provide fold change (FC) estimates on the log_2_ scale. The p-values from the Wald test were corrected for multiple testing using the Benjamini-Hochberg method to control the false discovery rate (FDR). Differentially expressed mRNAs in EF vs. PF were identified based on the criteria of |log_2_FC| >1 and adjusted p-value <0.05. The regularized log-transformed counts were used for downstream analyses such as hierarchical clustering and co-expression network analysis.

#### Protein significance analysis

Label-free quantification (LFQ) proteomics dataset was preprocessed with MaxQuant V2.6.7.0^68,69^ and previously published^10^. The quantification results were log-transformed (base 2) and normalized by equalizing the median intensities across MS runs. The mean normalized log_2_-intensities for EF and PF were compared and evaluated using two-sample t-test, followed by the Benjamini-Hochberg method for multiple testing correction.

#### Weighted gene co-expression network analysis

To evaluate co-expression patterns in RNA-seq (n=17,003 mRNA) and proteomics (n=1,134 proteins) datasets, a weighted gene co-expression network analysis (WGCNA V1.73)^70^ was performed. WGCNA uses pairwise correlation and hierarchical clustering to identify co-expression modules that are then related to the phenotype of EF vs. PF. Regularized log-transformed counts (RNA-seq) and normalized log_2_-intensities (proteomics) were used as the input to WGCNA. Soft-power thresholds to achieve the criterion of scale-free topology (R^2^>0.8) were determined for each dataset individually by running the pickSoftThreshold function. Then, automated network construction was performed with the parameters TOMtype=”signed”, mergeCutHeight=0.2, minModuleSize=20, and maxBlockSize=20,000. Co-expression modules were characterized based on the associations of their eigengene profiles with EF, evaluated using t-test followed by multiple testing corrections. GO enrichment analysis was performed for significant EF-associated modules (adjusted p<0.05). To evaluate potential enrichment of DE genes and proteins in the significant co-expression modules, the R package GeneOverlap V.1.44.0 was used to assess their overlaps based on Fisher’s exact test.

#### Directional integration of RNA-seq and proteomics data

To combine evidence of alcohol-induced changes in mRNA expression level and protein abundance, the differential analyses with the RNA-seq and LFQ proteomics datasets were integrated for the 1069 mRNAs/proteins quantified in both experiments. We used the directional p-value merging (DPM) method^22^, an extension of the empirical Brown’s method^71,72^ implemented in the R package ActivePathways V.2.0.5^73^, to construct a gene-level integration based on the p-values and fold changes reported from both analyses for each gene. The DPM method prioritizes genes in directional agreement while penalizes those in directional conflict.

#### Estimation and testing for protein turnover

To assess protein turnover, the previously reported kinetic data^10^ with 18 mice (n=9 EF and n=9 PF) were included in the integrative analysis. For each peptide, a time course of the total ^2^H-labeling was constructed. The kinetic data for native, unmodified peptides were used to evaluate the effect of alcohol on native protein turnover rate. Alcohol-induced change in turnover rate for acetylated protein was assessed in a site-specific manner, where each acetylated lysine site was assessed individually with only the acetylated peptides covering that site. The kinetic data were analyzed using a non-linear regression approach, where changes in turnover rates were assessed using an F-test^10^. As in the RNA-seq and LFQ proteomics analyses, turnover rate analyses conducted multiple testing correction, using the Benjamini-Hochberg procedure to control the FDR. A turnover rate change with an adjusted p<0.05 was considered statistically significant.

#### Functional enrichment analysis of deregulated genes and proteins

To evaluate functional enrichment of deregulated genes and proteins, enrichment analyses of differentially expressed genes and proteins with differential abundance or turnover rates were performed for Gene Ontology Biological Processes (GO:BP)^74^ and KEGG pathways^75,76^, using the R package clusterProfiler V.4.16.0^77–79^. Significant enrichments (FDR-adjusted p<0.05) were extracted for further visualization and interpretation. Top enriched terms were visualized in a dot plot or organized in an enriched map using the R package enrichplot V1.28.2. To compare enrichment results across different analyses, the significantly enriched GO:BP terms were visualized and summarized in Cytoscape^80^, using the EnrichmentMap pipeline^81,82^ that consists of the EnrichmentMap^82^, AutoAnnotate^83^, WordCloud^84^, and clusterMaker2^85^ applications. In the pipeline, EnrichmentMap visualized the enrichments as a network, where each node is a pathway and an edge represents the crosstalk between the pair of linked pathways in terms of the number of shared genes/proteins. Similar biological processes and functions were clustered using clusterMaker2, and the resulting clusters were labeled by AutoAnnotate based on word frequencies and order of the pathway descriptions. The nodes were rearranged for better visibility, and the cluster labels were manually curated to finalize the enrichment map.

#### Transcription factor enrichment analysis

For transcription factor enrichment analysis, proteins showing significant changes in abundance based on directional p-value merging or in turnover rates were used as input to the ChEA3 platform (https://maayanlab.cloud/chea3/)^26^, and queried against the Literature ChIP-seq TF-target gene set library.

#### Protein network and protein-metabolite analysis

Proteins showing significant changes in abundance or turnover rates were organized into a protein-protein interaction (PPI) network using the STRING database V.12.0^86^, with a stringent confidence score (>0.7). The STRING network was constructed in Cytoscape^80^ through the stringApp 2.0 application^24,87^ for further analysis and visualization. Network clustering was performed using the Markov clustering (MCL) algorithm, implemented in clusterMaker2^85^. Functional enrichment analysis was conducted for each of the top clusters, by querying gene sets from GO Biological Process (GO:BP), GO Cellular Compartment (GO:CC), Reactome pathways^88^ and KEGG pathways. After removing redundant terms using the default cutoff of 0.5, the most significant terms (up to the top four) were used to annotate the proteins within each cluster. Protein-metabolite analysis was also performed using stringApp 2.0^24^, in which metabolites of interests were queried against the STITCH database^36^ to identify their interacting proteins, which were subsequently filtered for liver tissue specificity using the TISSUES resource^36^.

## Data availability

Raw and processed RNA-seq data was deposited in the NCBI BioProject database and are publicly available under the accession number PRJNA1466568.

Mass spectrometry data was deposited in the ProteomeXchange Consortium under project accession number PXD055349.

All data supporting the findings of this study are included in the article and its Supplementary Information. Numerical source data underlying the graphs are provided in the Supplementary Data files. Additional data supporting the findings of this study are available from the corresponding author upon reasonable request.

## Author contributions

Conceptualization: T.K., T.-H.T., and L.E.N. Methodology: T.-H.T., M.A., S.I., M.R.M., U.S., S.G., W.C., and T.K., Formal Analysis: T.-H.T., T.K., and M.A., Visualization: T.-H.T., T.K., and M.A., Supervision: T.K., G.-F.Z., W.C., and L.E.N, Writing: T.K. and T.-H.T., Review and editing: T.-H.T., M.A., S.I., S.U., S.G., X.L.C., W.C., G.-F.Z., L.E.N., and T.K.

## Funding

This work was supported in part by the UH-NEOMED Neurodegenerative Disease Research Center (T.K.) and by National Institutes of Health grants R21AA029784 (T.K.), R21AG085590 (T.K.), R01AA030026 (X.L.C. and G.-F.Z.), and P50AA024333 (L.E.N.).

## Declaration of competing interest

The authors declare no competing interests.

## Legends of Supplementary Figures

**Figure S1.**
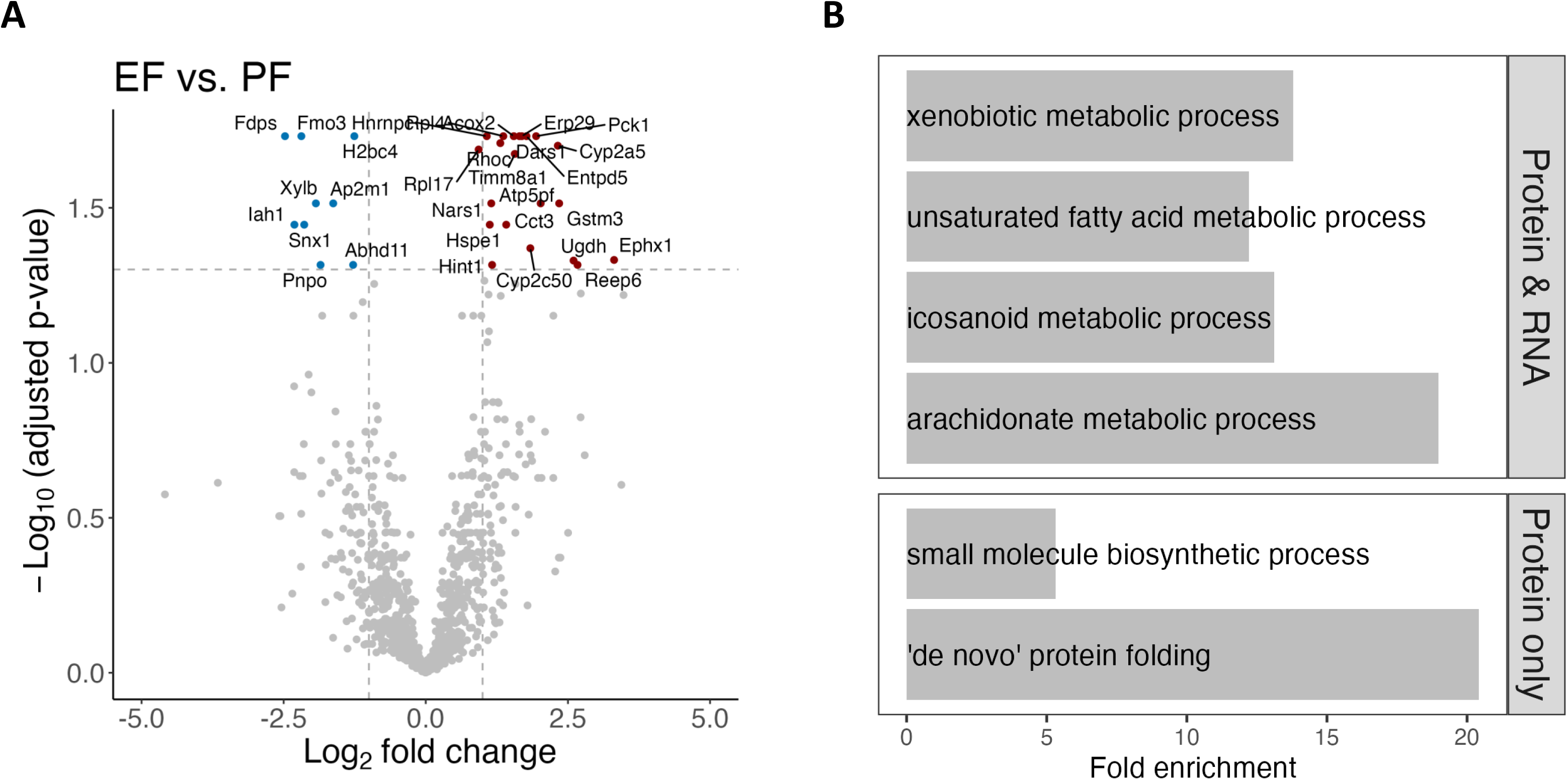
Reanalysis of LFQ proteomics data from Aghayev et al., 2025^10^. (**A**) Volcano plot summarizing the analysis of relative protein quantification. Of 1355 proteins quantified in 18 mouse livers (9 EF vs. 9 PF), 30 were differentially abundant (21 up-regulated and 9 down-regulated) with an adjusted p<0.05. (**B**) Eight significantly enriched GO:BP terms (**Supplementary Table S2**) associated with the 30 differentially abundant proteins (3 unique and 5 common with RNA-seq analysis). Redundant terms (‘de novo’ post-translational protein folding and long-chain fatty acid metabolic process) are not included.

**Figure S2.**
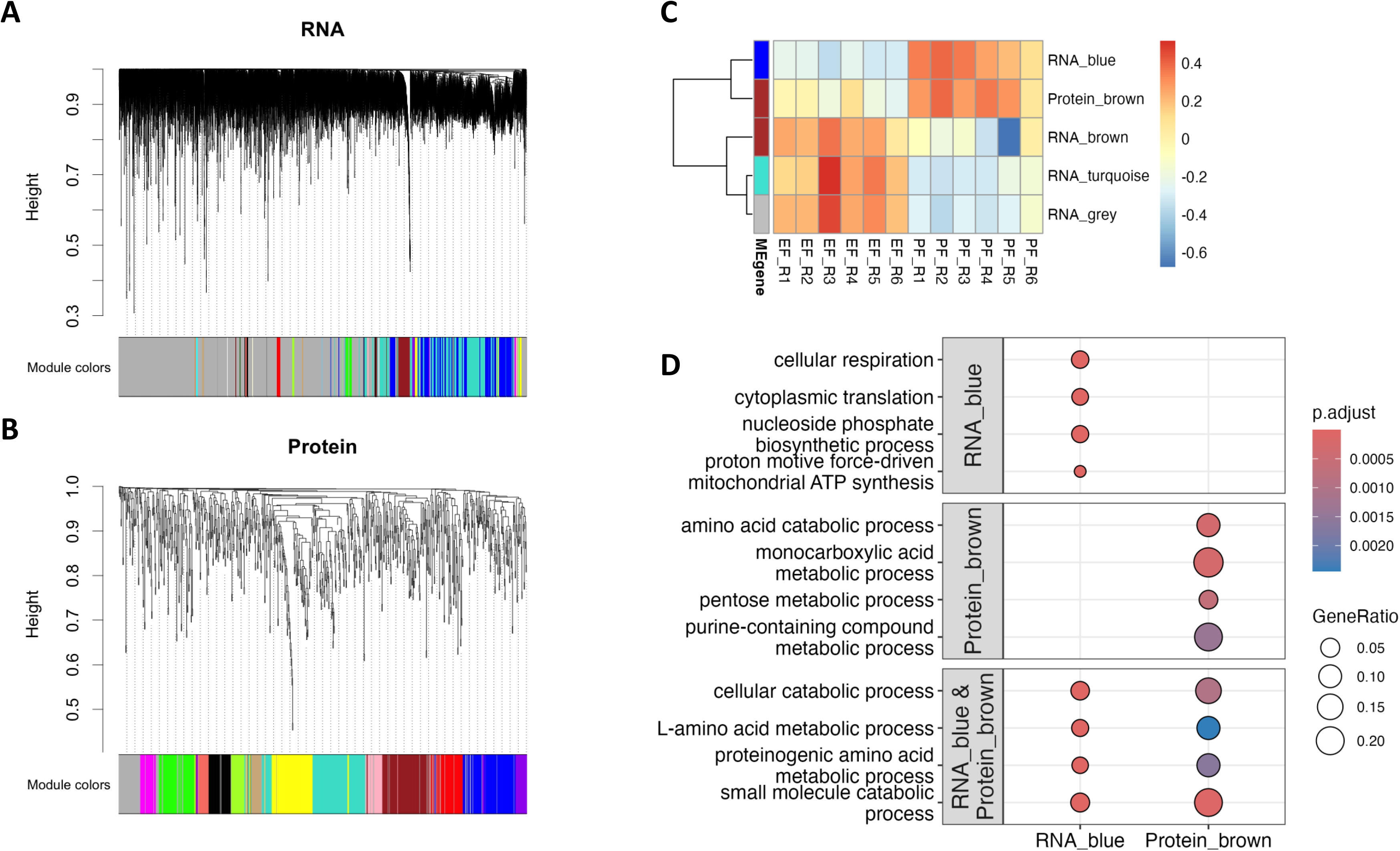
Co-expression network analysis of mRNAs and proteins. (**A**) Co-expression gene modules (n=30) identified from the RNA-seq dataset. (**B**) Co-expression protein modules (n=14) identified from the LFQ proteomics dataset. Module relevance was assessed by comparing module eigengenes, defined as the first principal component representing the overall expression profile of each module, between PF and EF groups. Among the identified modules, the mRNA “grey” (n=9890), “blue” (n=2150), “turquoise” (n=2615), and “brown” (n=528) modules, as well as the protein “brown” module (n=95), were significantly associated with EF (adjusted p<0.05) (**C**). (**D**) GO enrichment analysis of the RNA blue and protein brown co-expression modules, selected because the other significant modules represented broad biological themes not specifically linked to ALD. GO enrichment identified 101 and 27 biological processes associated with the mRNA blue module and protein brown module, respectively (**Supplementary Table S3**). Both modules were enriched for cellular catabolic and amino acid metabolic processes. The mRNA blue module showed additional enrichment for cellular respiration, cytoplasmic translation, nucleoside phosphate biosynthesis, and mitochondrial ATP synthesis, whereas the protein brown module was uniquely enriched for purine-containing compound, pentose, and monocarboxylic acid metabolic processes. To assess relationships between differentially expressed mRNA/proteins and co-expression modules within and across omics layers, deregulated features were compared with the RNA blue (n=2150) and protein brown (n=95) modules using the R package GeneOverlap. Consistent with reduced eigengene values in EF mice, downregulated mRNAs were significantly enriched in the mRNA blue module (138 overlapping genes; Fisher’s exact test, p=6.1×10^−88^), and downregulated proteins were enriched in the protein brown module (6 of 9 proteins; p=2.3×10^−10^). Upregulated mRNAs and proteins were also enriched in the mRNA blue (p=3.4×10^−29^) and protein brown (p=0.027) modules, respectively. No significant enrichment was observed across omics layers.

**Figure S3.**
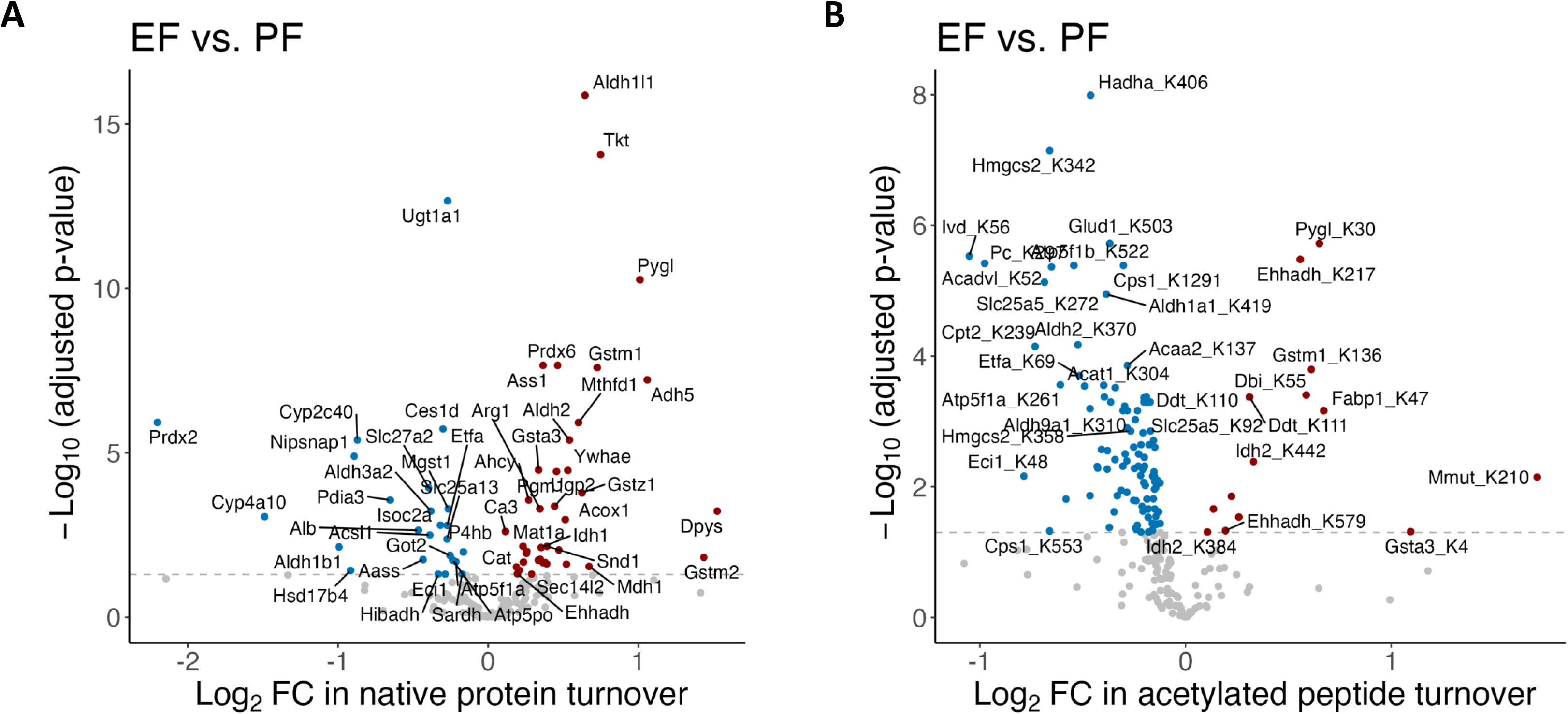
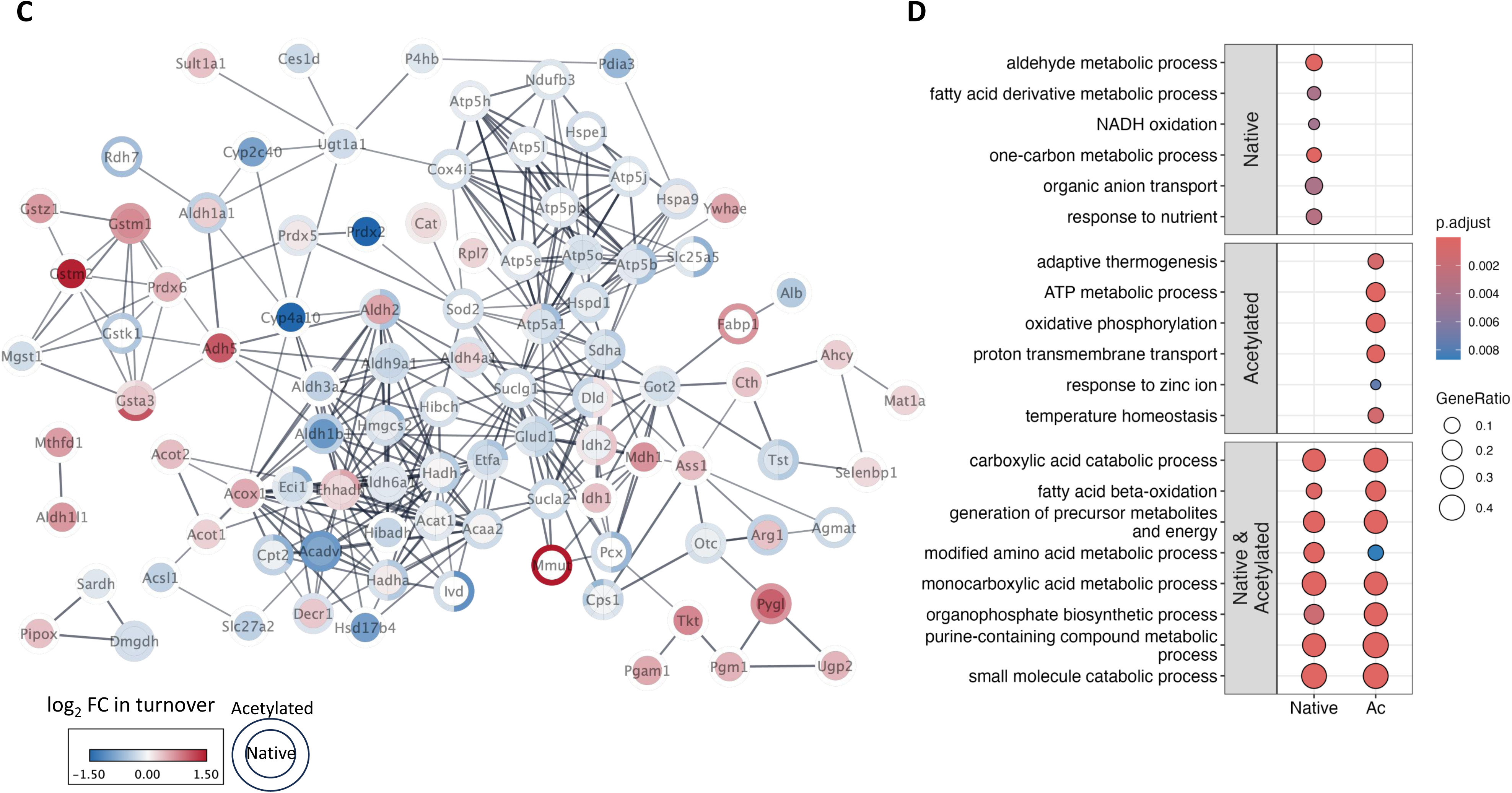
Reanalysis of kinetic proteomics data from Aghayev et al., 2025^10^. Alcohol significantly altered turnover rates of 63 native proteins (**A**) and 116 acetylated peptides corresponding to 67 proteins (**B**). (**C**) STRING interaction network of proteins exhibiting altered stability in either native or acetylated forms. Each circle represents a protein, colored based on its turnover rate changes in native (center) and acetylated (outer ring) forms. Red and blue colors correspond to increased and decreased turnover rates, respectively. Changes in protein stability were generally consistent among interacting neighboring proteins, suggesting coordinated regulation of protein turnover within functional pathways. (**D**) Representative GO:BP terms among 136 and 132 processes overrepresented in the 63 native proteins and 67 acetylated proteins, respectively (**Supplementary Table S5**). Of these, 66 GO:BP terms were shared between native and acetylated proteins with altered turnover rates, including amino acid, monocarboxylic acid, and sulfur compound metabolism; carboxylic acid and small molecule catabolic processes; and energy-generating pathways such as fatty acid beta-oxidation and organophosphate biosynthesis. In addition, 70 and 66 GO:BP terms were uniquely enriched among proteins with altered stability in native and acetylated forms, respectively. Representative native specific terms included aldehyde metabolism, fatty acid derivative metabolism, one-carbon metabolism, NADH oxidation, and organic anion transport, whereas acetylation-specific terms included adaptive thermogenesis, ATP metabolism, oxidative phosphorylation, proton transmembrane transport, and temperature homeostasis.

**Figure S4.**
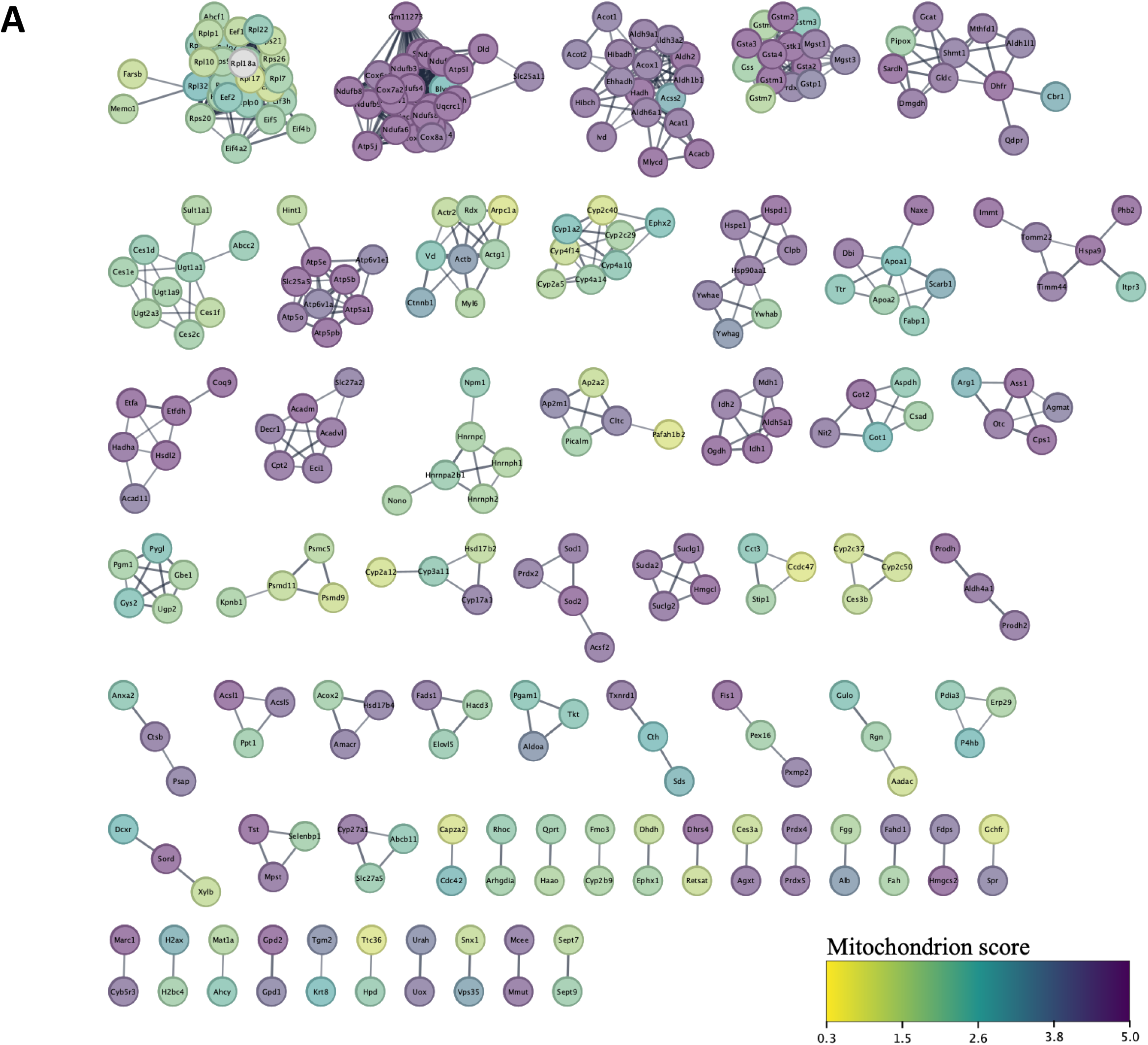
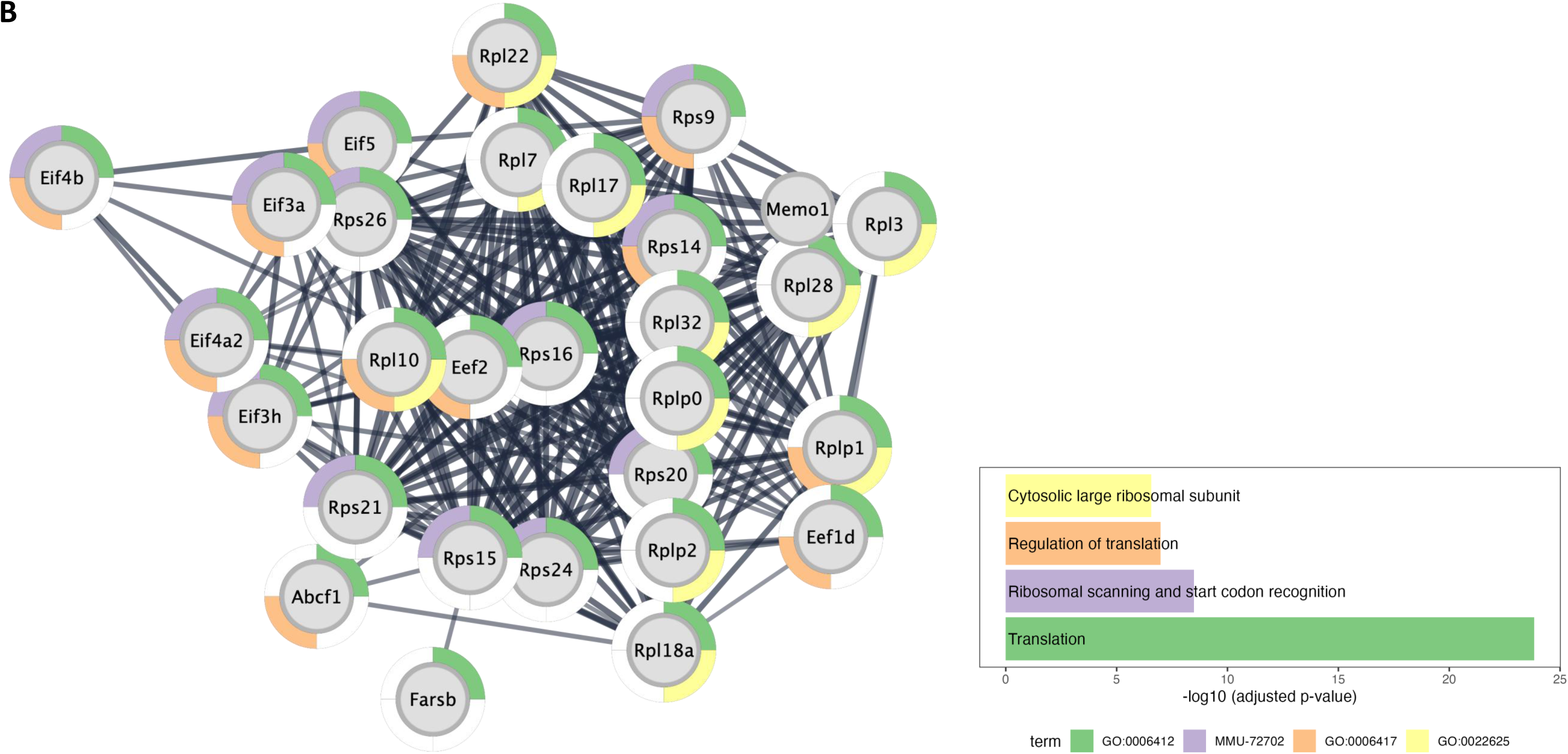
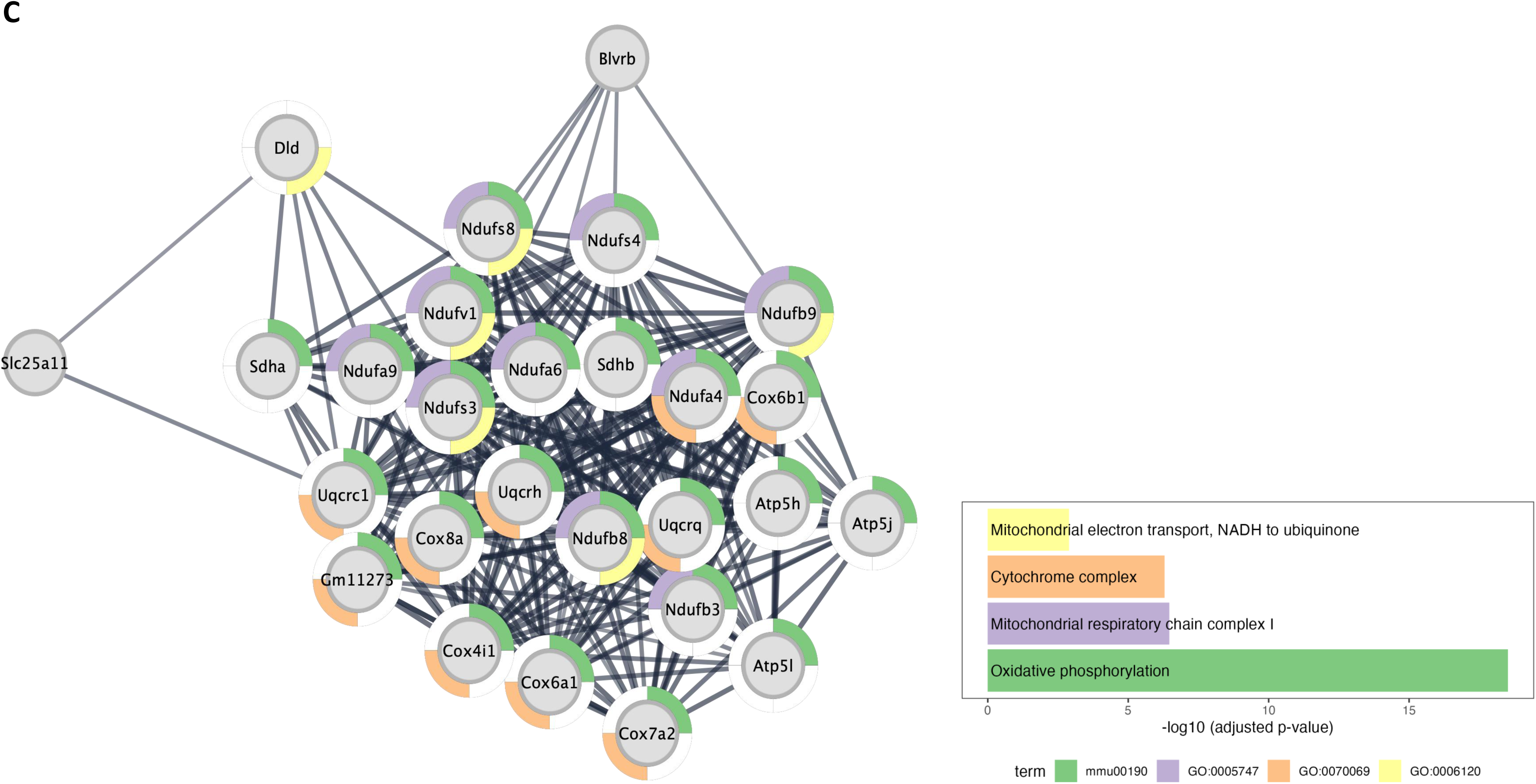
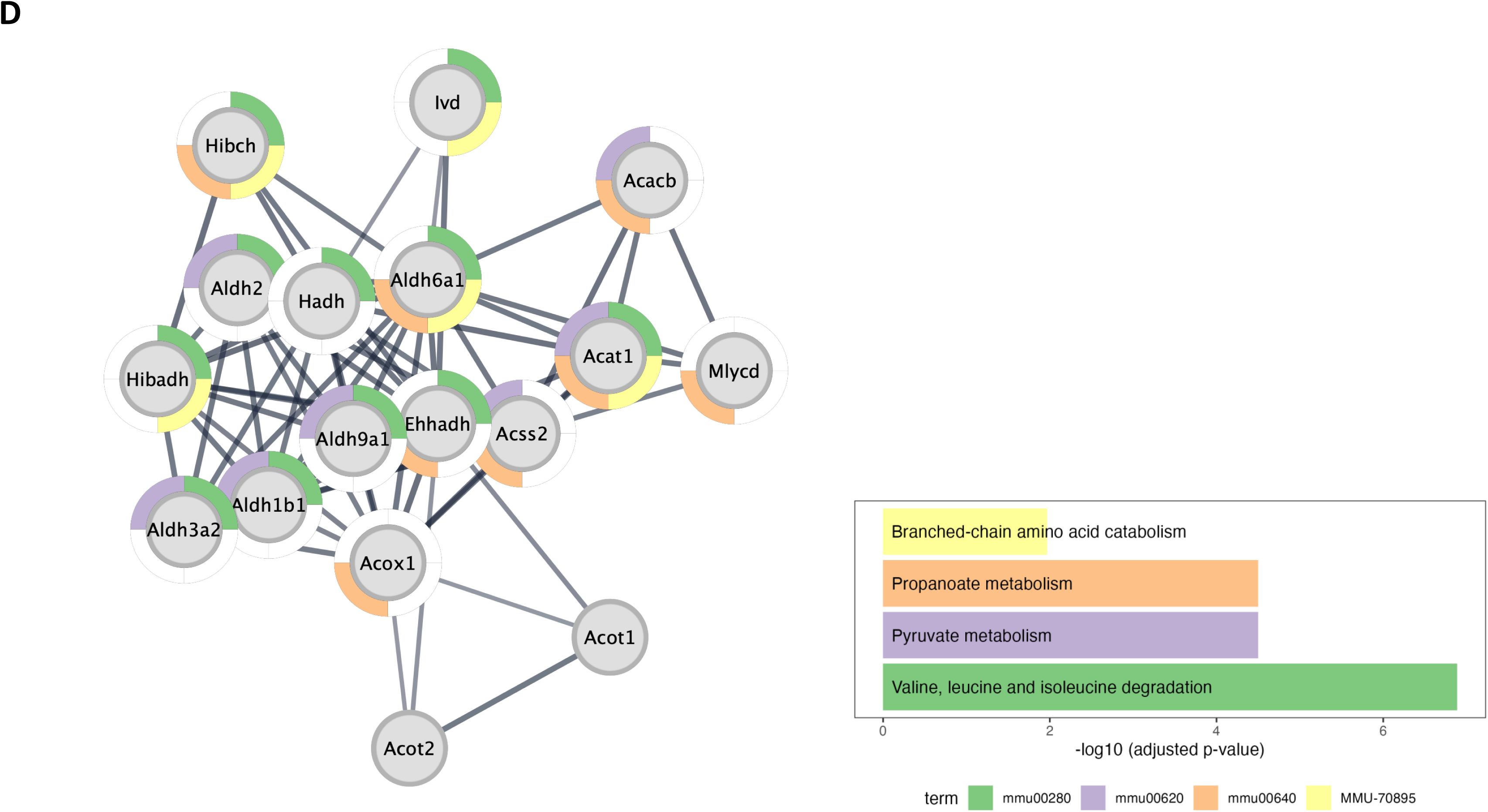
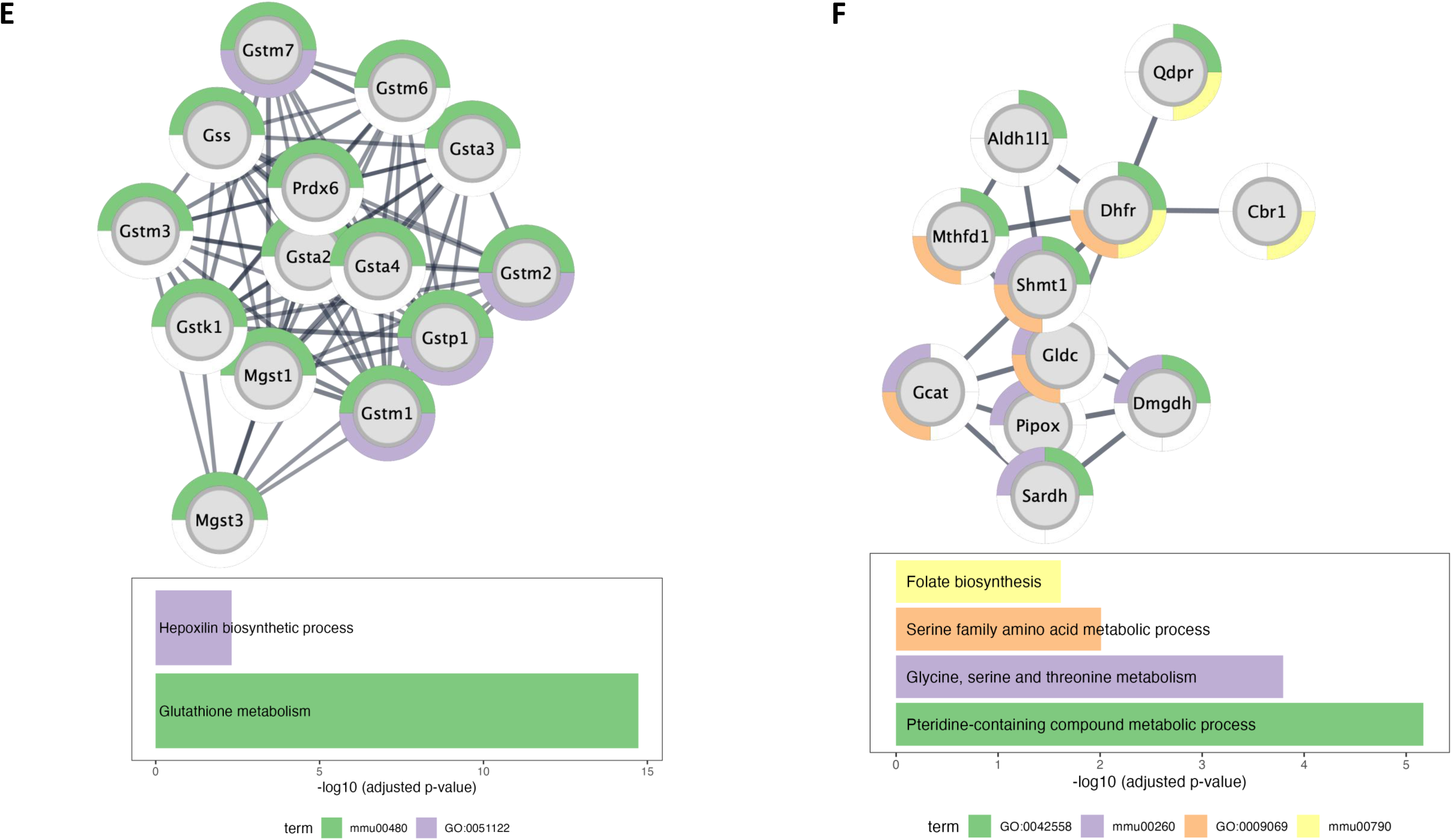
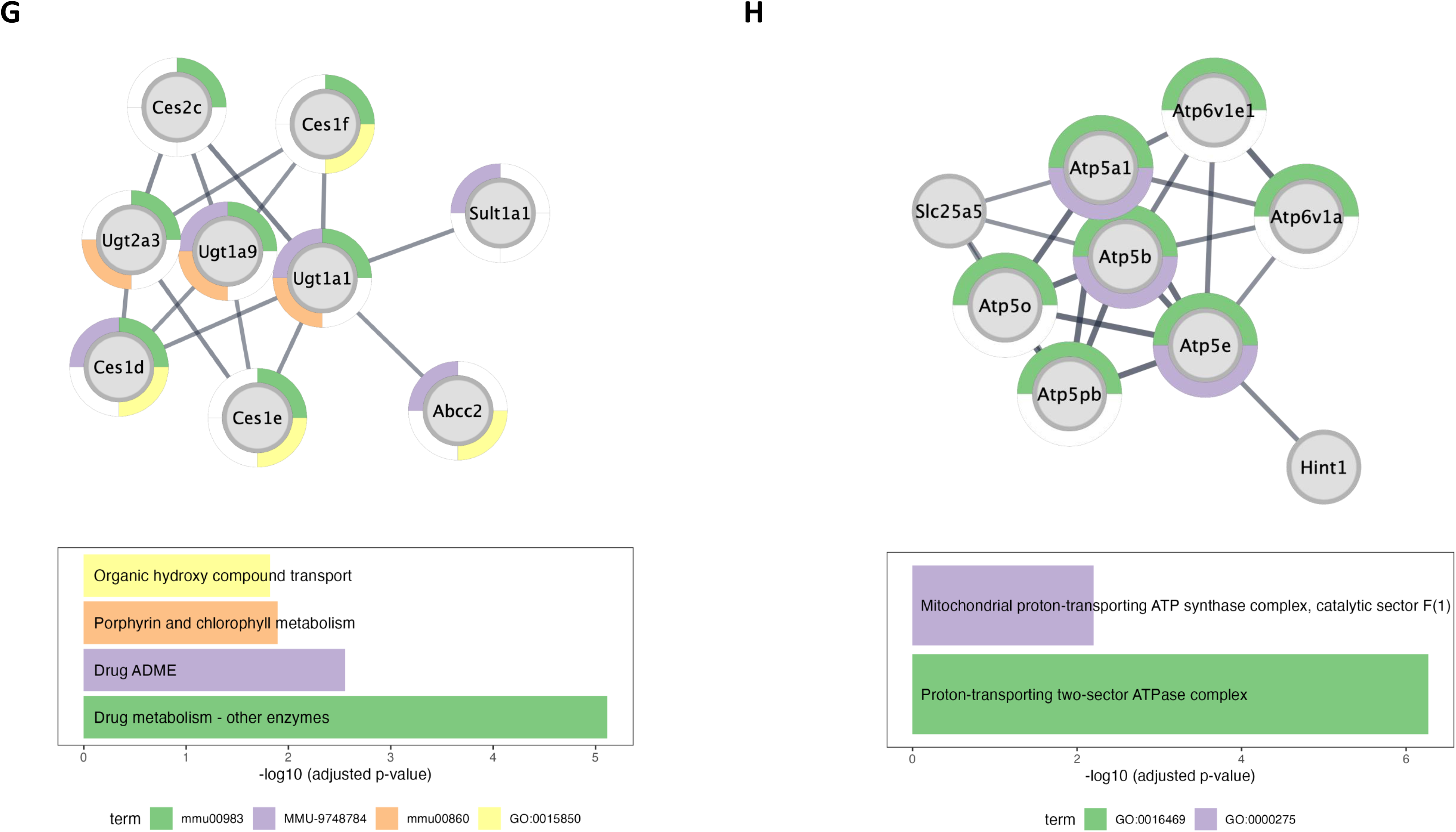
Interaction network of proteins with altered expression or turnover. (**A**) Clustered interaction network of proteins with a significant change in either expression (using the combined evidence) or turnover rate (native or acetylated). Each dot is a protein, colored based on its mitochondrion score. The clusters are ordered (from top to bottom, left to right) based on their size. Several clusters (e.g., #2-5, #7) are associated with mitochondrial proteins. Singleton proteins are not shown. (**B-H**) Subnetworks for Clusters #1-7 in (**A**), where the outer ring of each protein is annotated with the biological processes and molecular pathways significantly enriched in the cluster, compared to the entire protein interaction network.

**Figure S5.**
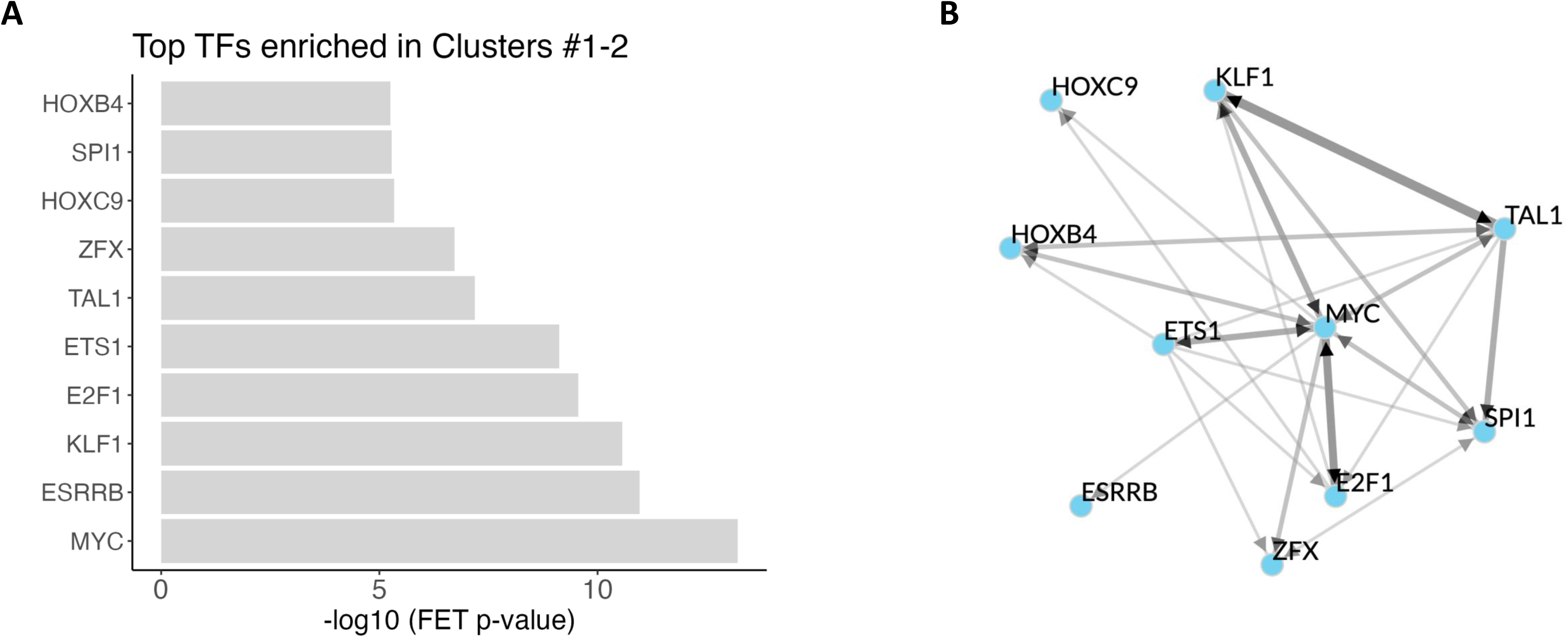
Transcription factors prioritized by ChEA3 for proteins with altered expression or turnover in Clusters #1-2. (**A**) Top-ranked transcription factors (ranks 1-10). (**B**) Interactions among the transcription factors shown in (**A**), based on evidence from ChEA3 libraries and, where applicable, supported by ChIP-seq data for directed interactions.

**Figure S6.**
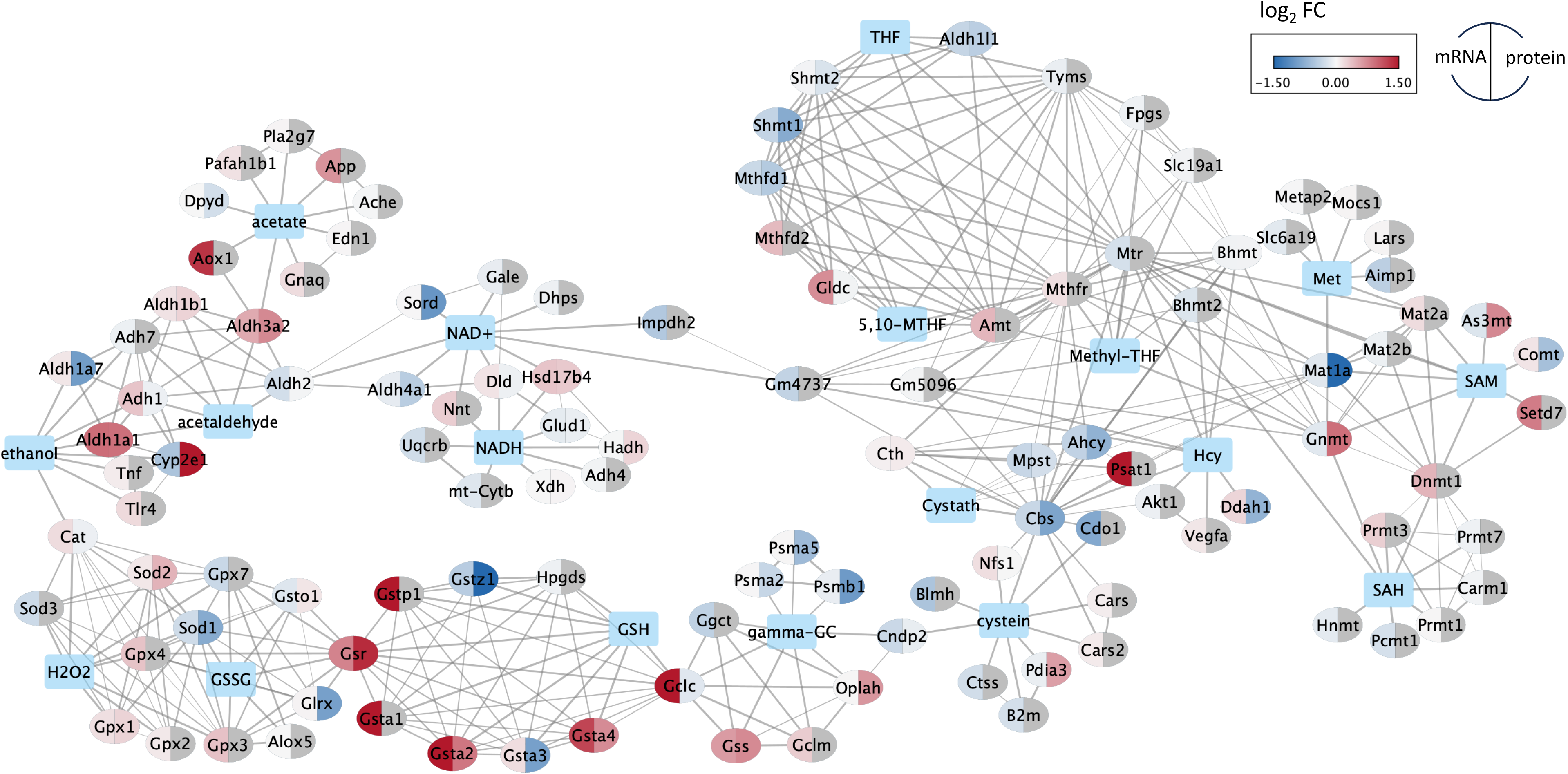
Protein-metabolite interaction network based on the STRING and STITCH databases, centered on key metabolites from the ethanol metabolism, methionine cycle and related pathways, transsulfuration, glutathione biosynthesis and utilization.

**Figure S7.**
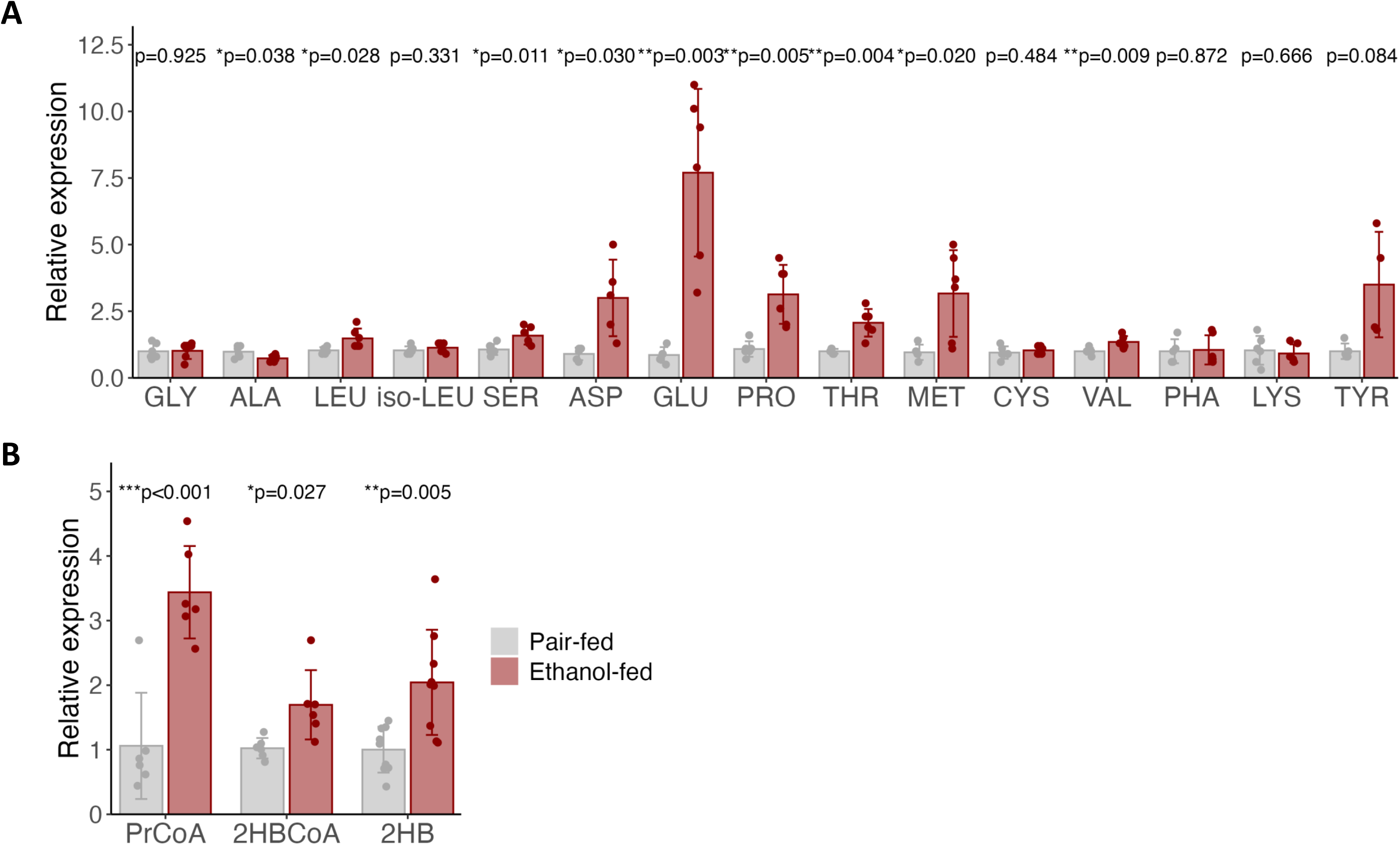
Reanalysis of metabolomics data from Aghayev et al., 2025^10^. Chronic alcohol consumption remodels hepatic amino acid metabolism and transsulfuration pathways. (**A**) Relative abundance of hepatic amino acids following chronic alcohol exposure. Elevated levels of multiple proteogenic amino acids are consistent with reduced urea cycle flux. In contrast, cysteine and glycine remain unchanged, supporting their preferential utilization for glutathione (GSH) synthesis in response to ethanol-induced oxidative stress. **(B)** Quantitative analysis of methionine cycle and transsulfuration intermediates. Chronic alcohol consumption increases 2-hydroxybutyrate, 2-hydroxybutyryl-CoA, and propionyl-CoA levels, consistent with enhanced transsulfuration flux to sustain antioxidant capacity.

## Notes

### Competing Interest Statement

The authors have declared no competing interest.

